# Role of RALBP1 in Oxidative Stress and Mitochondrial Dysfunction in Alzheimer’s Disease

**DOI:** 10.1101/2021.09.20.461132

**Authors:** Sanjay Awasthi, Ashley Hindle, Neha A. Sawant, Mathew George, Murali Vijayan, Sudhir Kshirsagar, Hallie Morton, Lloyd E. Bunquin, Philip T. Palade, J. Josh Lawrence, Hafiz Khan, Chhanda Bose, P. Hemachandra Reddy, Sharda P. Singh

## Abstract

The purpose of our study is to understand the role of the Ralbp1 gene in oxidative stress (OS), mitochondrial dysfunction and cognition in Alzheimer’s disease (AD) pathogenesis. The Ralbp1 gene encodes the 76 kDa protein Rlip (*aka* RLIP76). Previous studies have revealed its role in OS-related cancer. However, Rlip is transcriptionally regulated by *EP300*, a CREB-binding protein that is important for synaptic plasticity in the brain. Rlip functions as a stress-responsive/protective transporter of glutathione conjugates (GS-E) and xenobiotic toxins. OS causes rapid cellular accumulation of Rlip and its translocation from a tubulin-bound complex to the plasma membrane, mitochondria and nucleus. Therefore, Rlip may play an important role in maintaining cognitive function in the face of OS-related injury. This study is aimed to determine whether Rlip deficiency in mice is associated with AD-like cognitive and mitochondrial dysfunction. Brain tissue obtained from cohorts of wildtype and Rlip^+/-^ mice were analyzed for OS markers, expression of genes that regulate mitochondrial fission/fusion, and synaptic integrity. We also examined mitochondrial ultrastructure in mouse brains obtained from these mice and further analyzed the impact of Rlip deficiency on gene networks of AD, aging, inhibition of stress-activated gene expression, mitochondrial function, and CREB signaling. Our studies revealed a significant increase in the levels of OS markers and alterations in the expression of genes and proteins involved in mitochondrial biogenesis, dynamics and synapses in brain tissues of these mice. Furthermore, we compared the cognitive function of wildtype and Rlip^+/-^ mice. Behavioral, basic motor and sensory function tests in Rlip^+/-^ mice revealed cognitive decline, similar to AD. Gene network analysis indicated dysregulation of stress-activated gene expression, mitochondrial function, and CREB signaling genes in the Rlip^+/-^ mouse liver. Our results suggest that the Rlip deficiency-associated increase in OS and mitochondrial dysfunction could contribute to the development of OS-related AD processes. Therefore, the restoration of Rlip activity and endogenous cytoprotective mechanisms by pharmacological interventions is a novel approach to protect against AD.

## Introduction

Alzheimer’s disease (AD), the most common form of dementia, is irreversible, progressive and heterogeneous, having several etiologies and pathological processes that evolve as it progresses from mild cognitive impairment to dementia (1). AD is the 6th leading cause of death and is the only disease among the 10 leading causes of death in the United States that cannot be cured, prevented or slowed. AD occurs in two forms, early onset familial and late-onset sporadic. Synaptic damage and loss correlate strongly with cognitive function in patients with AD (2-5). Several years of research revealed multiple cellular changes are involved in disease progression, including microRNA deregulation, synaptic damage, the proliferation of astrocytes & glia, oxidative stress, mitochondrial structural & functional abnormalities, hormonal imbalance, formation and accumulation of Aβ and p-Tau and neuronal loss (6, 7).

In AD, increased levels of free radicals and oxidative stress are extensively reported (8-13). Generation of superoxide anion (O_2,_^-•^, the unpaired ‘radical’ electron shown as ^•^) reacts with polyunsaturated fatty acid (PUFA) to yield toxic free radicals (i.e. OH^•^, O_2_^-•^ etc.) and electrophilic lipid metabolites (LOO^•^). Environmental toxins, heavy metals, cigarette smoke, chronic inflammation and most other risk factors for AD have in common the ability to initiate or promote lipid peroxidation (LPO) (8, 14). Given its high lipid composition, it is not surprising that LPO is highly toxic to brain tissue and is a hallmark of late-stage AD and other neurodegenerative diseases (14). No approved drugs dissolve free-radical generating AD plaques or tangles and no chemical antioxidants are sufficiently effective free radical scavengers to terminate LPO.

Developing drugs that activate the expression of glutathione (GSH)-linked antioxidant enzymes (GLAE), which metabolize LOO^•^ and electrophilic lipids, is critically important because impaired GLAE activation is a key pathogenic mechanism of AD (15, 16). GLAE is upregulated by NRF2, a transcription factor (TF) stabilized by reaction of electrophilic lipids with its binding partner, KEAP1. NRF2 binds an antioxidant response element (ARE), inducing transcription of GLAEs. Another stress-responsive TF CREB (cAMP regulatory element binding protein) along with its co-regulators CBP (CREB-binding protein) and EP300 (CBP-coactivator), as well as the master TF for heat-shock proteins, HSF and its co-activator p53 interact with and modulate NRF2 functions (17-19).

Mercapturic acid pathway transporters remove mercapturic acid precursor GSH-electrophile conjugates (GS-E) from cells (20-22). RALBP1 encoded RLIP76 (referred to here as Rlip), a 76 kDa splice variant protein that bound the clathrin-dependent endocytosis (CDE) complex through the clathrin adaptor protein AP2 (25, 26). Its ATPase activity and GS-E transport were coupled with the internalization of peptide hormone/receptor complexes by CDE (25, 26), and it was present in an intracellular tyrosine kinase signaling complex that also contains Grb2/Nck, and Shc (25, 27-29). Rlip knockout mice had impaired CDE, structurally abnormal mitochondria and high levels of OS (30-32). NRF2 and GLAE should be increased and GSH should be low due to increased consumption by GSH-utilizing enzymes. Surprisingly, both NRF2 and GLAE levels were lower than in wildtype mice and GSH levels were elevated, perhaps due to deficiency of GSH-consuming enzymes.

We recently reported that heterozygous Rlip deficiency exerted genome-wide alterations of gene promoter CpG island methylation (31). Gene expression array of Rlip^+/-^ mice confirmed that the top differentially expressed canonical pathways involved OS and endocytosis. NRF2 was among the top differentially upregulated genes by RNASeq studies and NRF2 gene promoter was among the top differentially methylated regions (DMR) (31). Our studies suggest a link between NRF2 and the curious finding that many of the top differentially expressed pathways were neuronal (CREB-signaling in neurons, axonal guidance and long-term potentiation).

In the present study, we report for the first time that Rlip^+/-^ mice had some neurocognitive deficits that resemble AD. Thus, the Rlip^+/-^ mice represent a unique model of AD in which NRF2-regulated antioxidant enzyme activities are lower in Rlip^+/-^ mice similarly, and some key mitochondrial proteins were similarly reduced. We further explored a novel possibility that the observed altered mitochondrial biogenesis, dynamics and synaptic proteins associated with the AD by Rlip. These studies will offer novel insight into the regulation of OS defenses in AD and can lead to new strategies for the treatment of AD and offer insights into the OS models of dementia.

## Results

### Rlip^+/-^ mice exhibit neurocognitive abnormalities that resemble an AD mouse model

Behavioral, basic motor and sensory function tests in mouse models can reveal behavioral impairments. Since OS markers are significantly higher in Rlip^+/-^ mice, we suspected that oxidative damage in the brain might lead to cognitive impairments in Rlip^+/-^ mice. Therefore, we tested Rlip^+/-^ mice in neurocognitive tests on 5 WT and 5 Rlip^+/-^ mice using Open Field Maze, RotaRod, Y Maze and Morris Water Maze tests. Open-field maze studies provided the first preliminary evidence to support the idea of 7-9 month-old mice having a potential AD or AD-like phenotype (**Figure 1A-B**). Rlip+/- mice were less mobile than WT mice (F(1,8)=7.4, *p*=0.0260). Rlip^+/-^ mice tended to exhibit a shallower turn angle was observed in comparison to WT, which trended towards significance for both turn angle and (F(1,8)=5.2, *p*=0.051), and, correcting for distance traveled, meander (F(1,8)=4.5, p=0.067) (**Figure 1A-B**). These findings were extended in additional studies showing that significant differences in distance traveled (*p*=0.034), mobility (*p*=0.0044), turn angle (*p*=0.025) and meander (*p*=0.0033) become more pronounced in the same mice with increased age (10-12 months), suggestive of accelerated aging in Rlip^+/-^ mice (**Figure 1C-D**). There was also a trend for Rlip^+/-^ mice to spend more time in the periphery than center than WT mice (*p*=0.0607), suggestive of an emerging anxiety phenotype.

**Figure 1.**
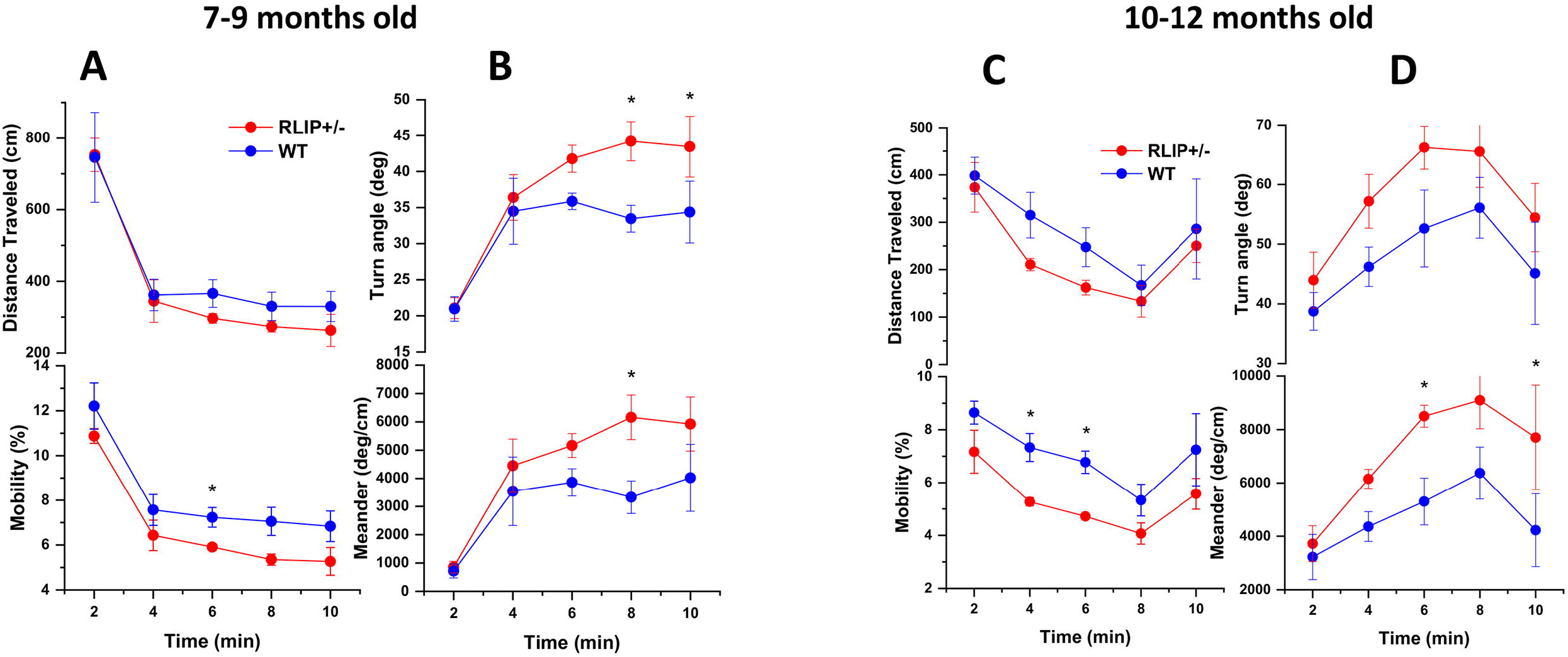
Aging increases phenotypic differences between Rlip^+/-^ and wildtype (WT) mice. (**A**) For a population of 5 Rlip het mice and 5 WT mice of 7-9 months, both groups exhibited habituation over the course of 10 minutes with reduced distance traveled (F(4,32)=30.8; *p*<0.0001) and mobility (F(2.2,17.9)=37.35;*p*<0.0001). Rlip+/- mice were less mobile than WT mice (F(1,8)=7.4, *p*=0.0260). (**B**) Rlip+/- mice tended to exhibit a shallower turn angle was observed in comparison to WT, which trended towards significance for both turn angle and (F(1,8)=5.2, *p*=0.051), and, correcting for distance traveled, meander (F(1,8)=4.5, *p*=0.067). (**C**) For a population of 5 Rlip +/- mice and 5 WT mice of 10-12 months, both groups exhibited habituation over the course of 10 minutes with reduced distance traveled (F(4,32)=7.62; *p*=0.0002) and mobility (F(4,32)=7.38;p=0.0003). (**D**) Rlip^+/-^ mice tended to exhibit a shallower turn angle compared to WT, which was more pronounced in aged mice (F(1,8)=8.12, *p*=0.022), and, correcting for distance traveled, meander (F(1,8)=17.0, p=0.0033). Significant differences (*p*<0.05) in specific time points from post-hoc analysis (Fisher’s LSD) are indicated with asterisks (*).

We then tested these mice in learning and memory tests. The Rotarod performance test was used to evaluate WT and Rlip^+/-^ mouse behavioral task performance, a natural fear of falling, motor coordination and fatigue. Mice were placed on a rod that rotates at accelerating speed (0 to 40 rpm) over the course of 5 minutes. As shown in **Figure 2A**, Rlip haploinsufficiency negatively affected the performance times of the mice. A two-tailed *t-test* revealed a significant drop in the performance time on the apparatus by Rlip^+/-^ mice (Mean=23.55 sec, *p* < 0.009), when compared with WT cohorts (Mean=74.85 sec, < 0.0022).

**Figure 2.**
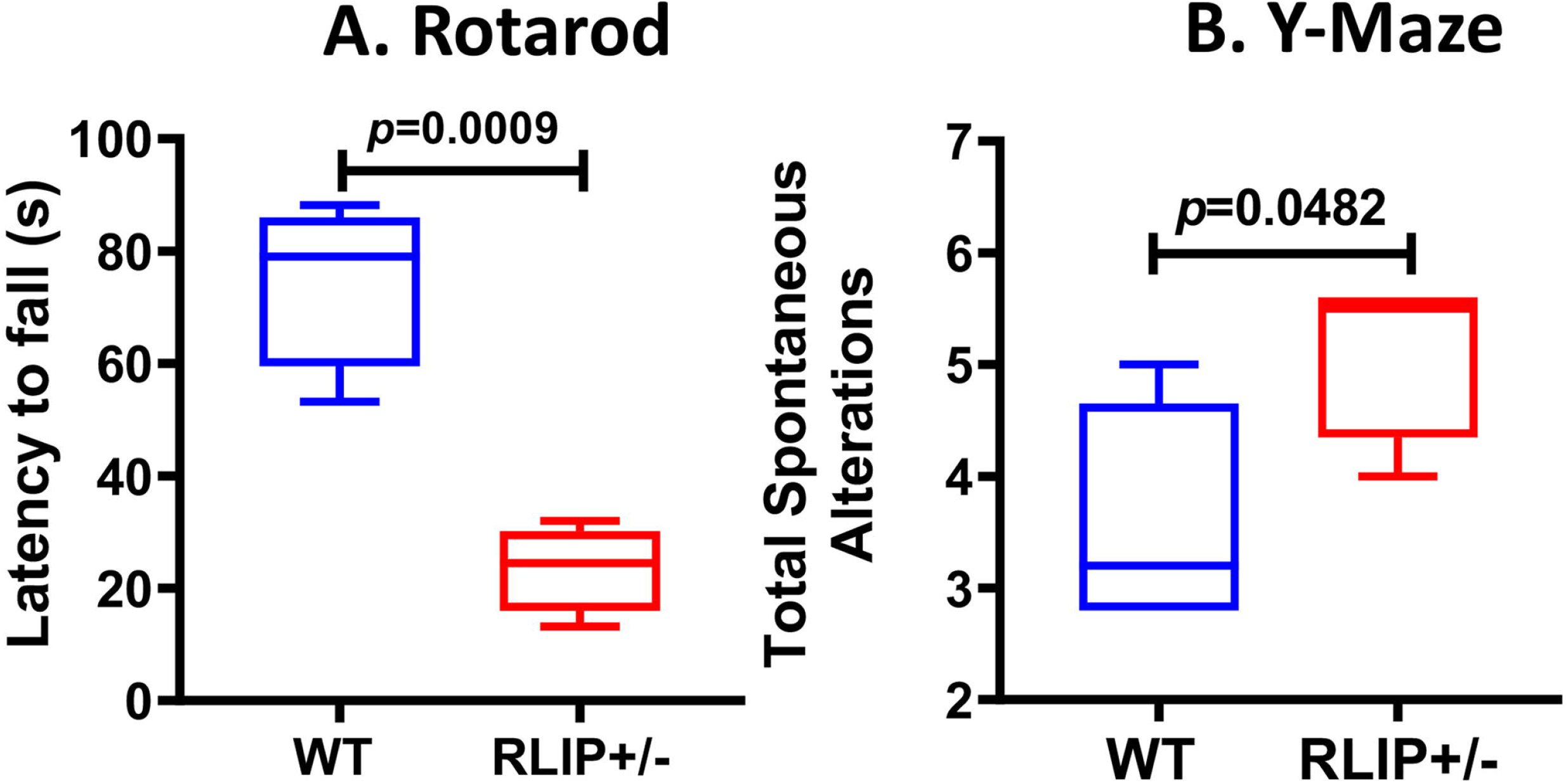
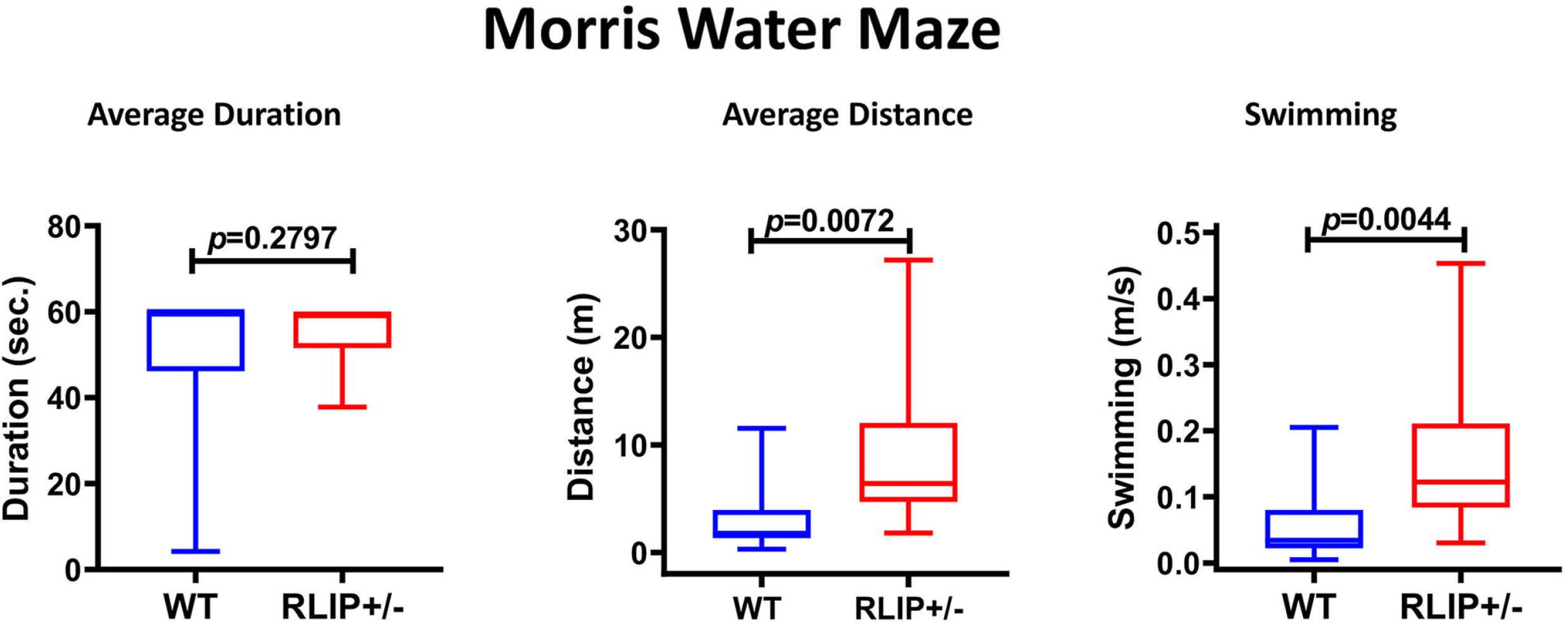
Rlip deficiency decreases the latency to fall of mice on Rotarod and increases total spontaneous alterations in Y-maze. Analysis of locomotor activity including **A**) Latencies to fall off an accelerating rotarod. **B**) spontaneous assessed in Y-maze test. 5 Rlip+/- mice and 5 WT mice of 10-12-month-old were assessed for neurological and behavioral phenotypes. Significant differences (p<0.05) from post-hoc analysis (Fisher’s LSD) are indicated in each panel.

The Y maze is a behavioral test used to measure the willingness of mice to explore novel environments, which in turn reveal any cognitive deficits in transgenic mice. Typically, in this test mice will investigate a new arm of the maze over returning to a previously visited arm. This involves many parts of a rodent’s brain including the prefrontal cortex, hippocampus, basal forebrain, and septum. Surprisingly, total spontaneous alternation is significantly higher in Rlip^+/-^ mice than in their wt counterparts (**Figure 2B)**. The Morris water maze test also tends in the same direction as the Y-maze test (**Figure 3**). Significantly increased average distance (*p*=0.0072) and swimming time (*p*=0.0044) were found for Rlip+/- mice compared to age-matched, WT mice, indicating that Rlip+/- mice showed impaired cognitive function **(Figure 3)**.

**Figure 3.**
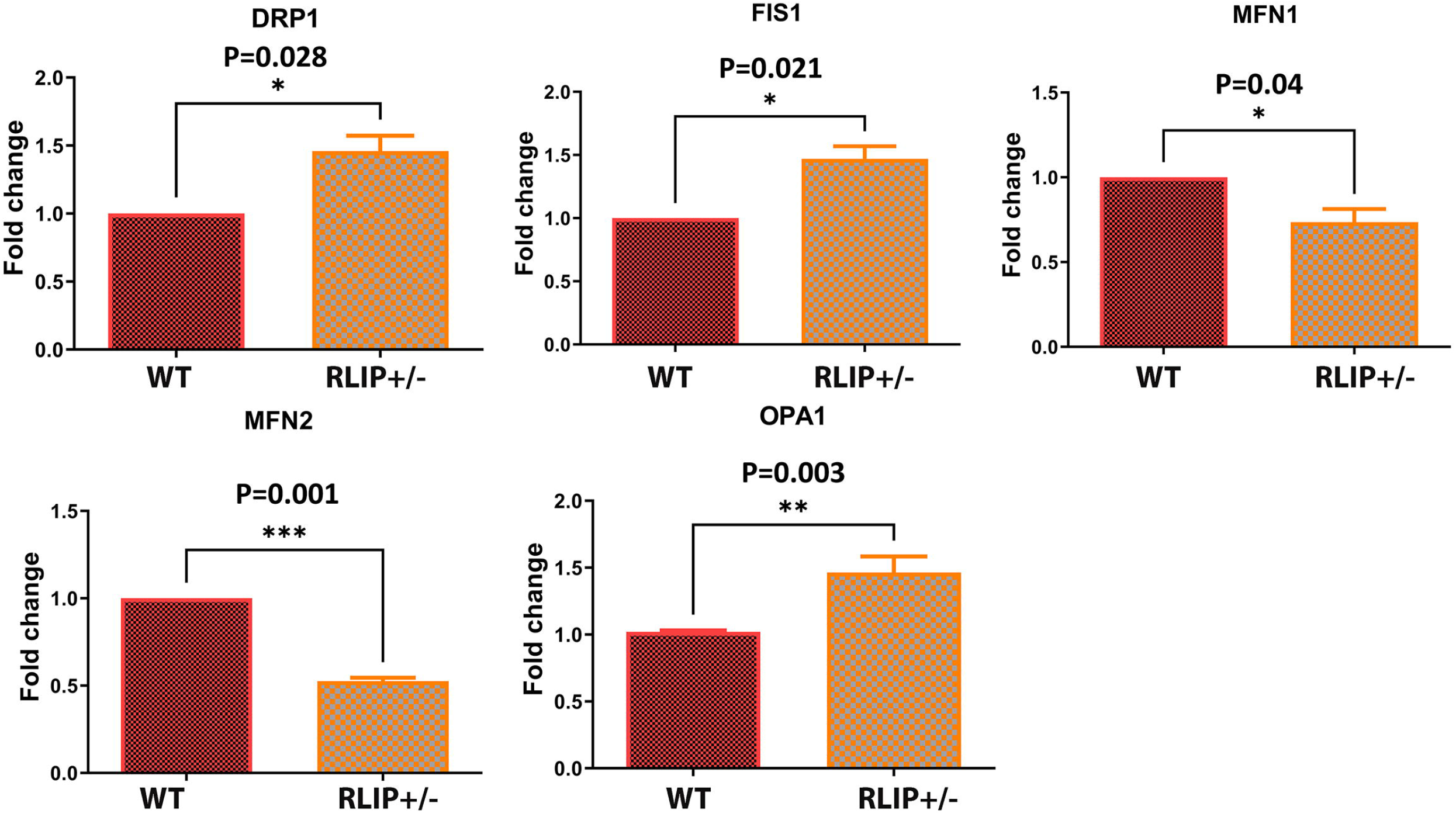
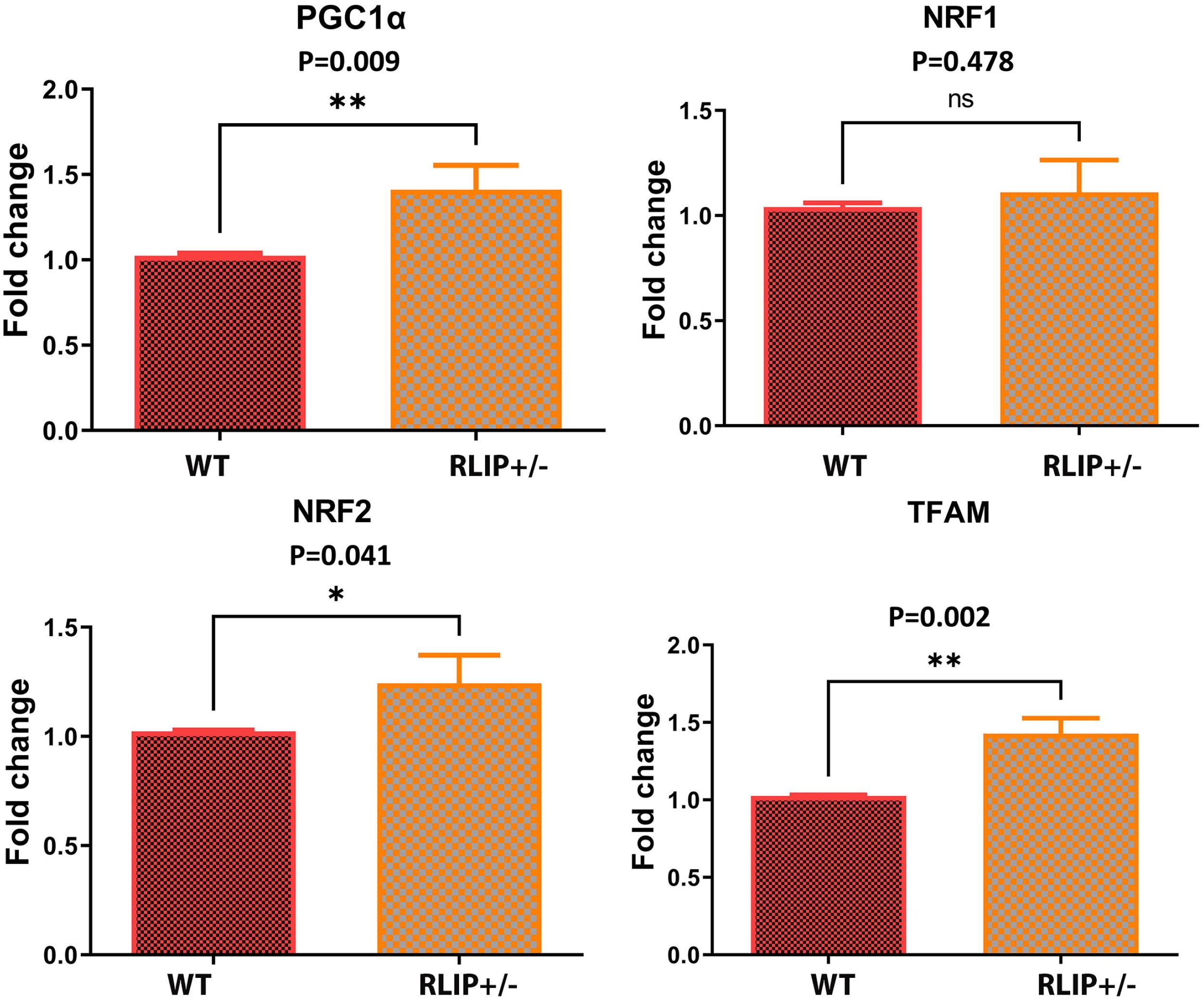
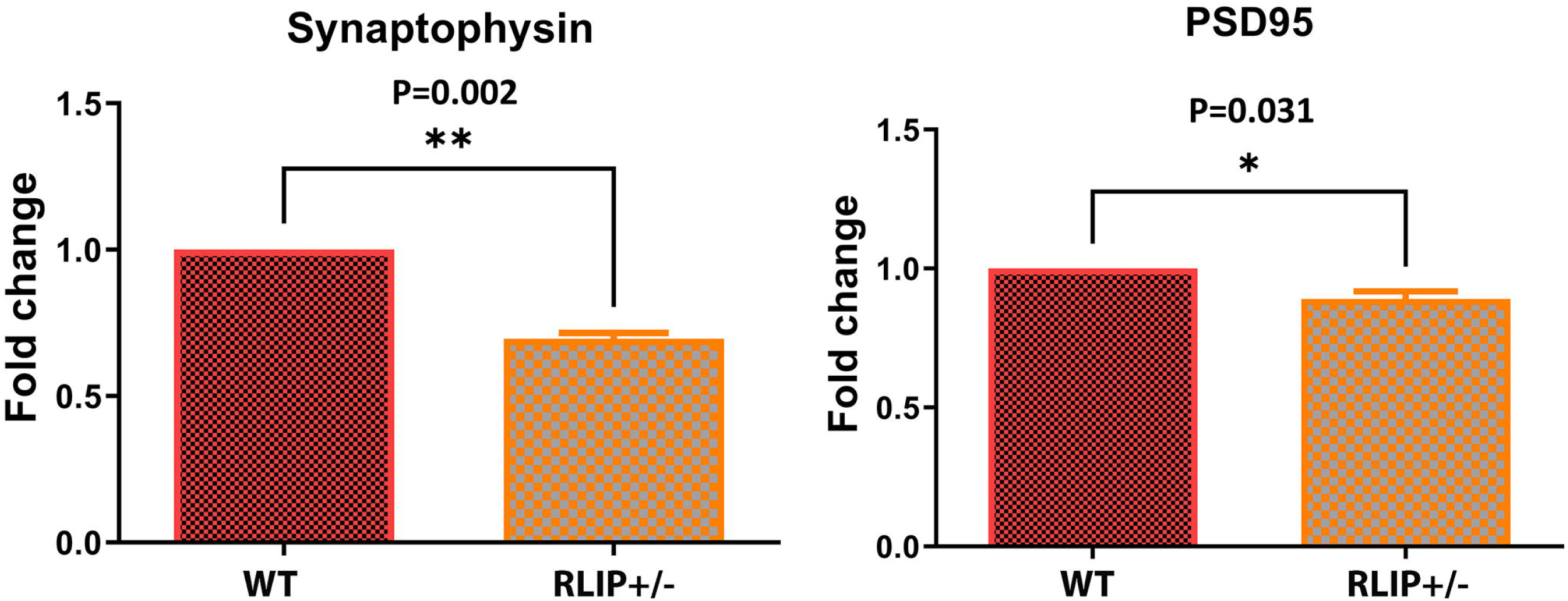
Rlip deficiency alters the willingness of mice for spatial learning. Spatial learning was assessed as a function of a training day with respect to the following parameters: Average Duration, Average Distance Traveled in the target quadrant, and Swimming speed in the target quadrant. Data are presented as the mean ± SD (n = 5 animals of each genotype) and significance was determined by post-hoc analysis (Fisher’s LSD) are indicated in each panel.

### Effect of Rlip Deficiency on Expression of Genes that Regulate Mitochondrial Fission/Fusion and Synaptic Function

#### Mitochondrial dynamics

Rlip deficiency impairs mitochondrial fission (32). To determine, whether, Rlip deficiency affects mitochondrial dynamics (fission and fusion), we measured mRNA levels of fission genes (Drp1 and Fis1) and fusion genes (Mfn2, Mfn2 and Opa1) in Rlip^+/-^ and aged-matched control WT mice. As shown in Figure 4A, Fission genes Drp1 ((*p*=0.028) and Fis1 ((*p*=0.021) were significantly increased in Rlip+/- mice relative to WT mice. On the other hand, mitochondrial fusion genes (Mfn1, *p*=0.040; Mfn2,*p*=0.001) were significantly reduced in Rlip+/- mice relative to WT mice. However, mRNA levels of Opa1 were increased in Rlip+/- mice. Overall, these observations strongly suggest that the presence of impaired mitochondrial dynamics in Rlip+/- mice, similar to AD cells (42,44), APP transgenic mice (33) and Tau transgenic mice (34).

**Figure 4.**
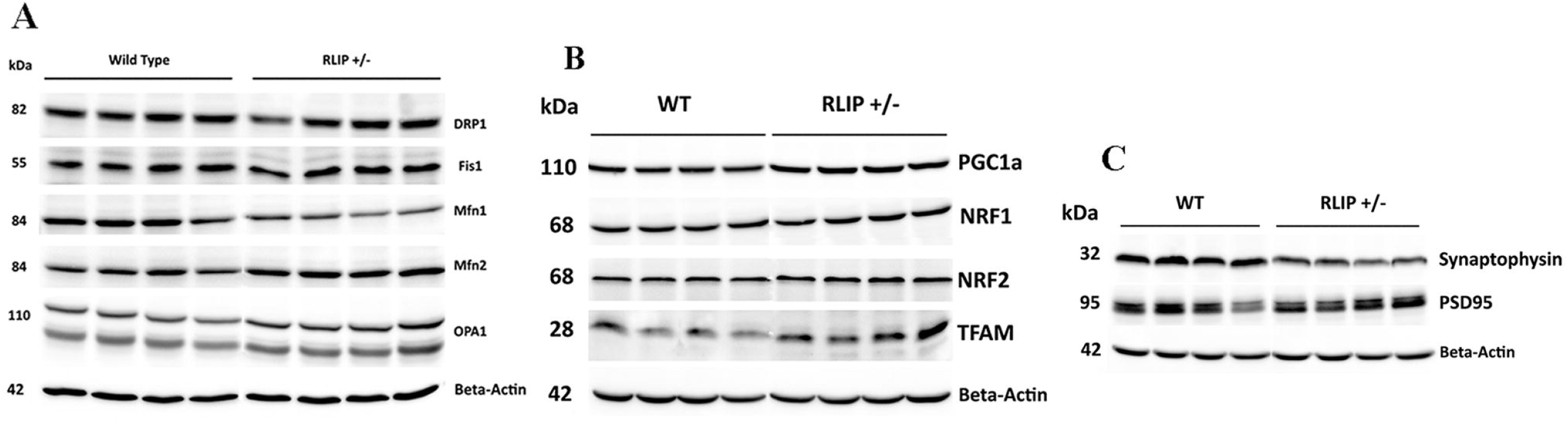
Effect of Rlip Deficiency on Expression of Genes that Regulate Mitochondrial Dynamics, Synaptic and Bioenergetics Function. qRT-PCR was performed by previously described methods on mouse brain homogenates from 3 WT and 3 Rlip^+/-^ mice. Levels of Mitochondrial function regulating transcripts encoding Dynamics (A), Synaptic (B) and Bioenergetics (C) protein were measured in brains of WT and Rlip+/- animals. Gene expression levels were normalized to the beta-actin transcript and calculated as described in Materials and methods. Asterisks denote statistically significant differences between WT and Rlip+/- groups of animals, compared pairwise Student’s t-tests.

#### Mitochondrial biogenesis

Mitochondrial biogenesis was assessed in Rlip+/- mice, in order to understand the impact of Rlip gene deficiency in mice. Interestingly, mitochondrial biogenesis genes (PGC1a, *p*=0.009); Nrf2 *p*=0.041; and TFAM, p=0.002) were increased in Rlip+/- mice relative to WT mice (**Figure 4B**). These observations indicate that increased mitochondrial biogenesis may be a compensatory response to reduced mitochondrial function in Rlip+/- mice.

#### Synaptic genes

As shown in **Figure 4C**, synaptic genes, synaptophysin (*p*=0.002) and PSD95 (*p*=0.031) were significantly reduced in Rlip+/- mice.

These observations indicate that mRNA levels of mitochondrial dynamics, biogenesis and synaptic genes were altered, very similar to mouse models of AD.

### Immunoblotting Analysis

To determine the effects of Rlip depletion on protein levels, we performed the immunoblotting analysis of protein lysates prepared from brain tissues of WT and Rlip^+/-^ mice.

#### Mitochondrial dynamics

As shown in **Figure 5**, we found increased in Fis1 (P=005) in the brain of the Rlip^+/-^ mouse when compared with the WT counterparts. Mitochondrial fusion proteins Mfn1 (*p*=0.033) and Mfn2 (*p*=0.005) were significantly reduced in Rlip+/- mice relative to WT mice.

**Figure 5.**
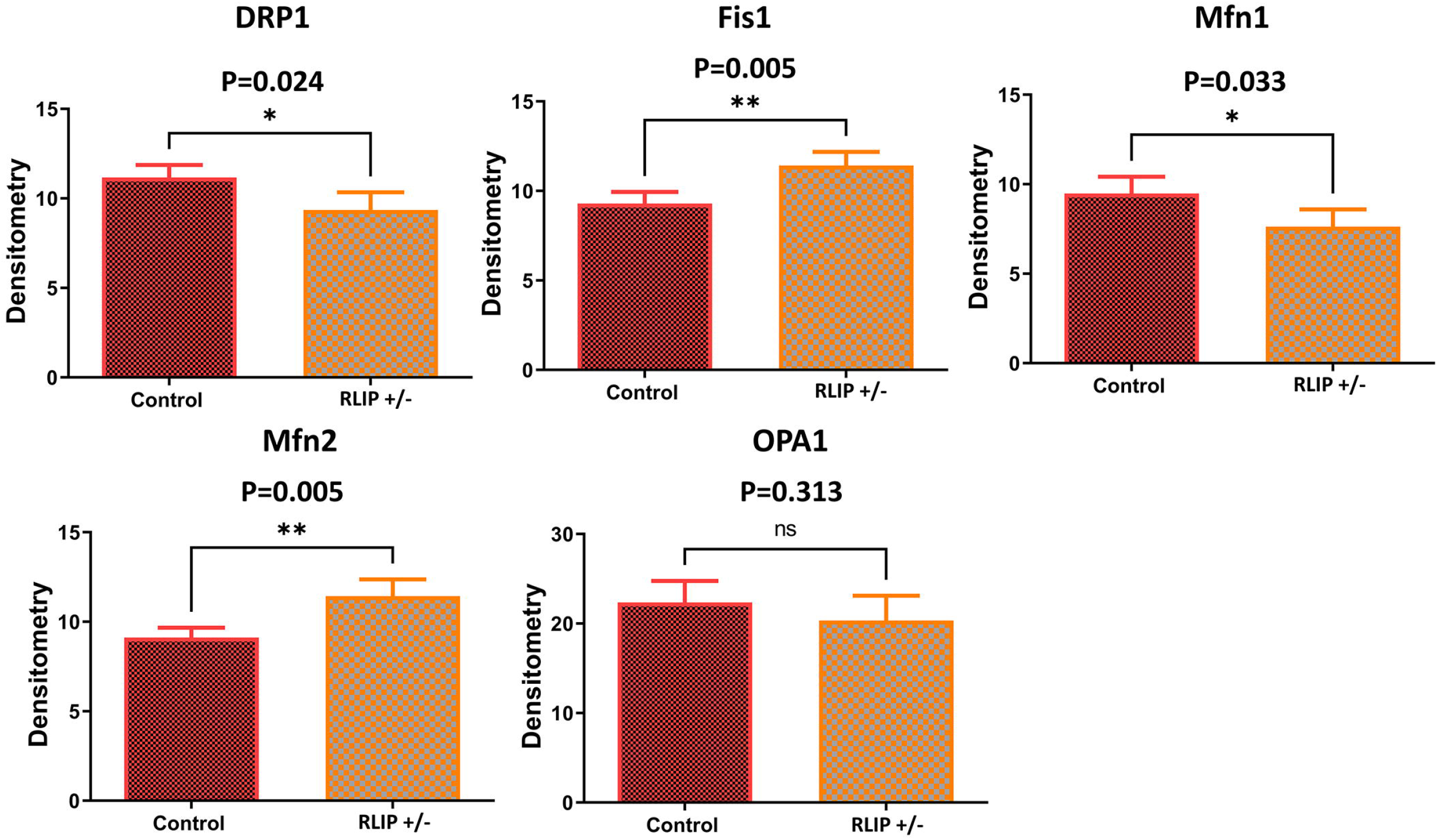
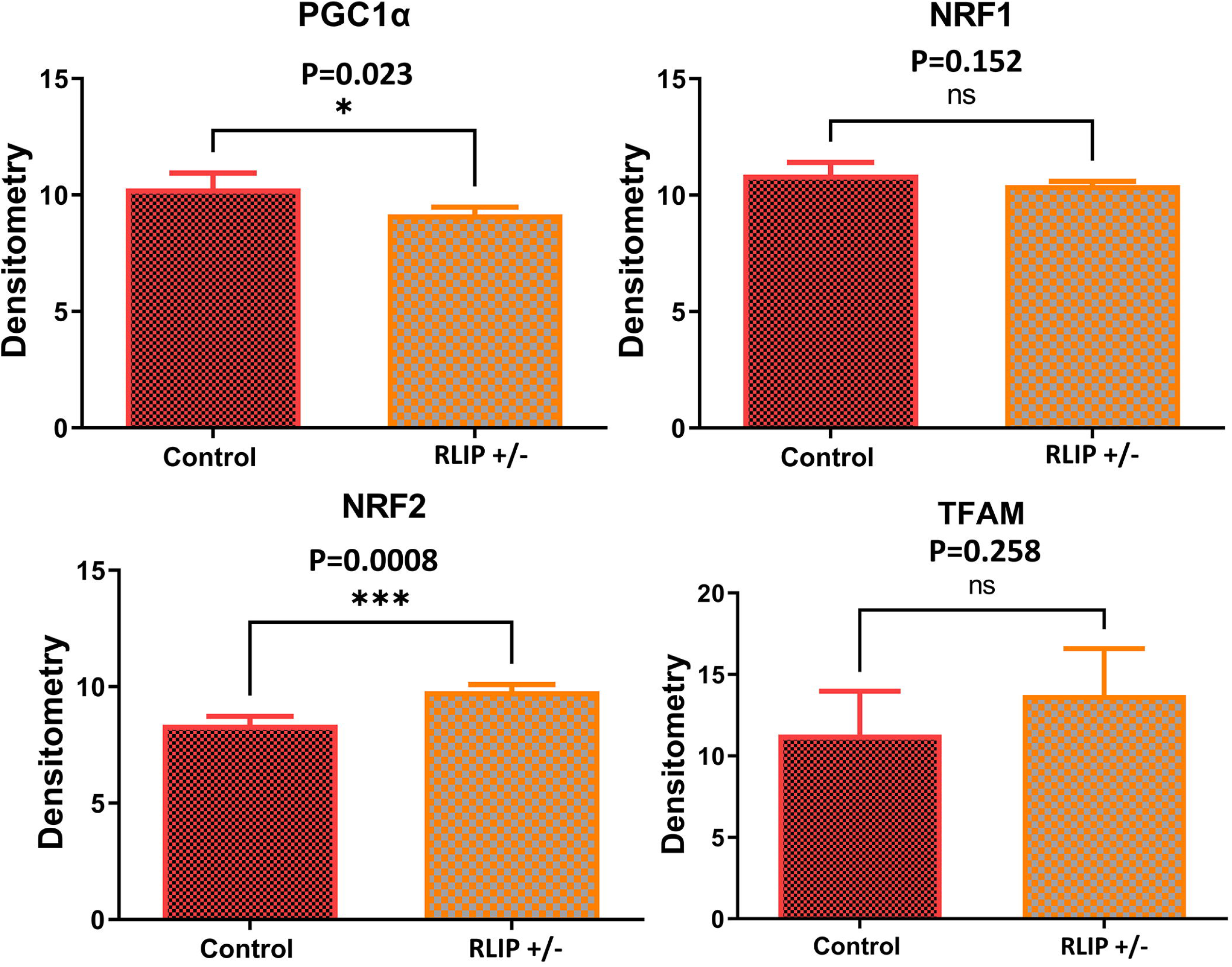
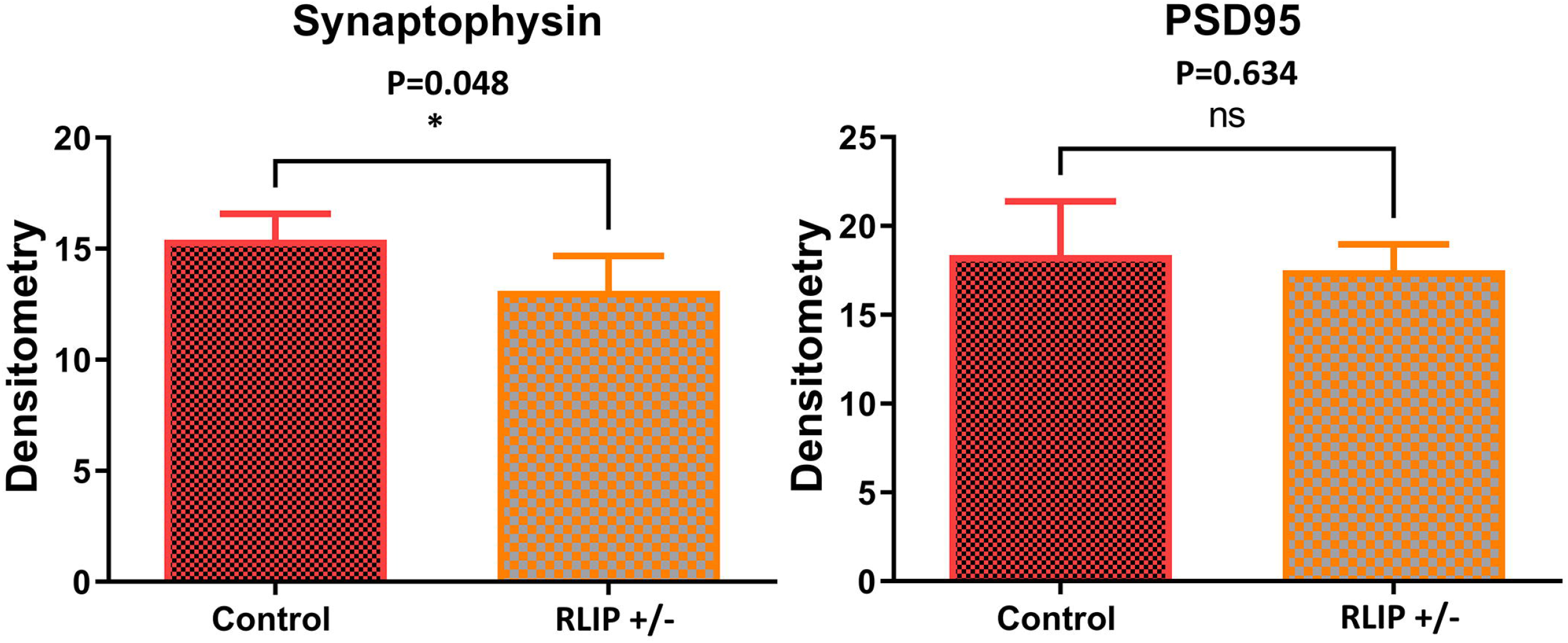
Expression of key mitochondrial proteins in the brain of WT and Rlip+/- mice. Rlip+/- deficiency alters proteins mitochondrial protein regulating Dynamics (A), Synaptic (B) and Bioenergetics (C) functions. Brain homogenates from WT and Rlip+/- mice were examined by Western blot analysis (**5 A-C**). Grayscale values of the protein bands, normalized with β-actin using ImageJ software (developed by NIH), are shown in **5 D-F** to indicate the relative expression levels of proteins in a representative Western blot. Asterisks denote statistically significant differences between WT and Rlip+/- groups of animals, compared pairwise Student’s t-tests.

#### Mitochondrial biogenesis

Significantly increased mitochondrial biogenesis proteins PGC1a, P=0.023; Nrf2, P=0.0008 were found in Rlip+/- mice relative to WT mice. However, protein levels of Nrf1 and TFAM did not change in Rlip+/- mice. These observations indicate that biogenesis is affected by reduced Rlip in mice.

#### Synaptic proteins

Similar to mRNA levels, synaptic proteins, synaptophysin (P=0.053) and PSD95 (P=0.043) were significantly reduced in Rlip+/- mice, when compared with WT mice, indicating that synaptic activity is defective in Rlip+/- mice, what was observed in APP and tau transgenic mice of AD (33, 34).

### Immunofluorescence Analysis

#### Mitochondrial dynamics proteins

We performed immunofluorescence of mitochondrial dynamics, biogenesis and synaptic proteins, mainly to understand the impact on AD-affected brain regions, hippocampus and cortex. As shown in **Figure 6A** the immunoreactivities of Drp1 (*p*=0.0210) and Fis1 (*p*=0.0018) were significantly increased in the Rlip^+/-^ mice relative to WT mice. On the other hand, fusion protein Mfn1 was significantly reduced in Rlip^+/-^ mice (*p*=0.217) relative to WT mice. Our immunofluorescence data concur with qRT-PCR (mRNA levels) and immunoblotting (protein levels) data.

**Figure 6.**
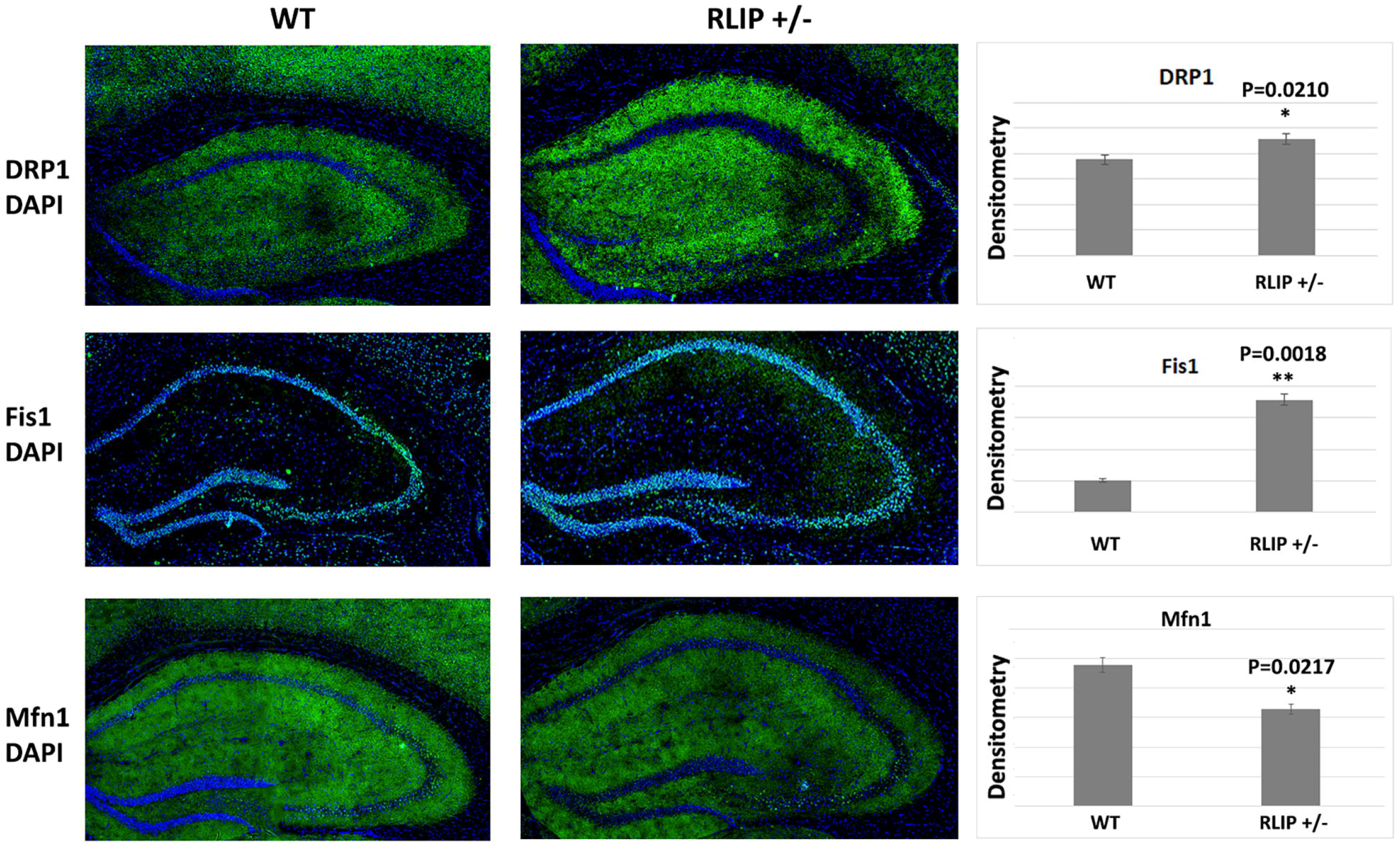
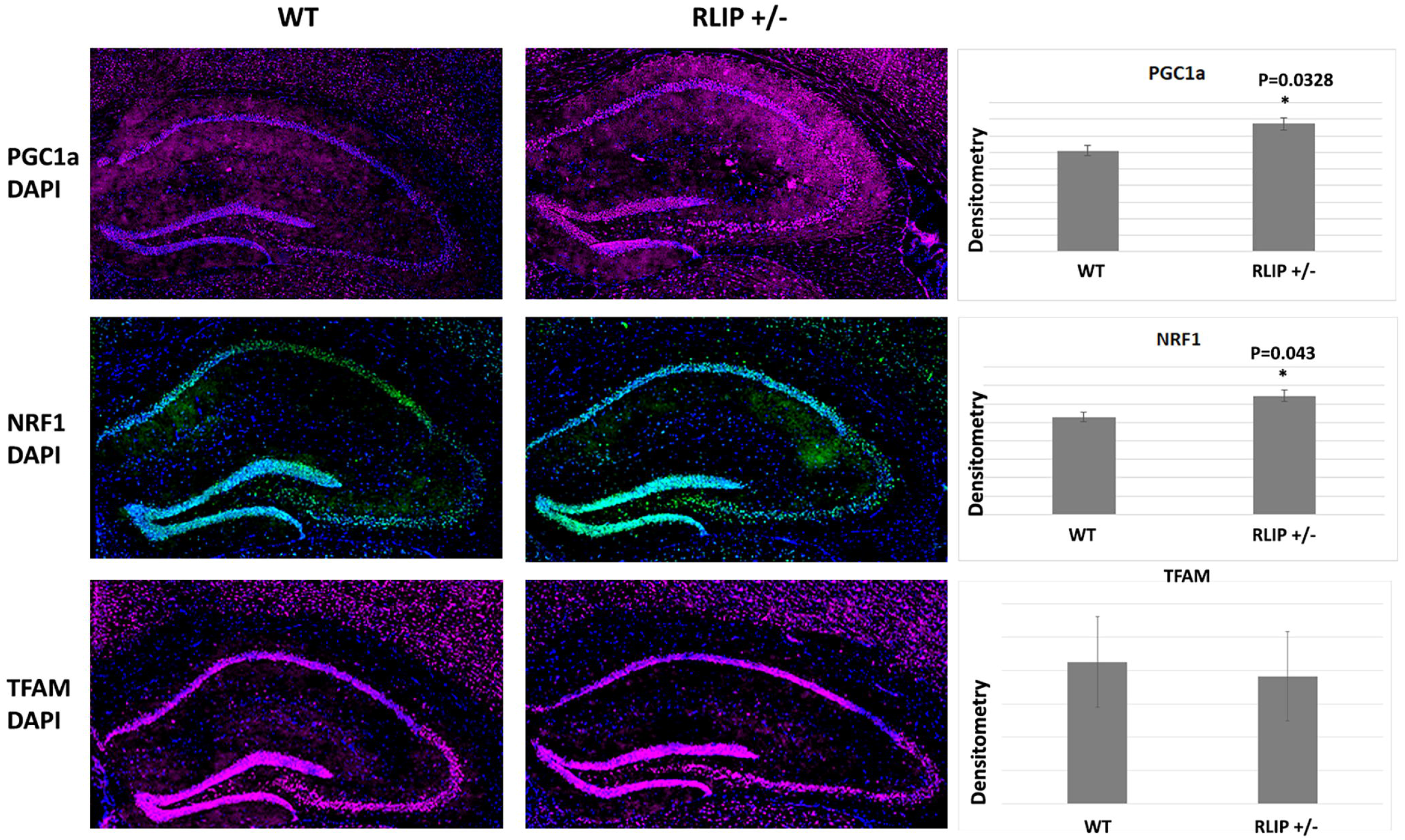
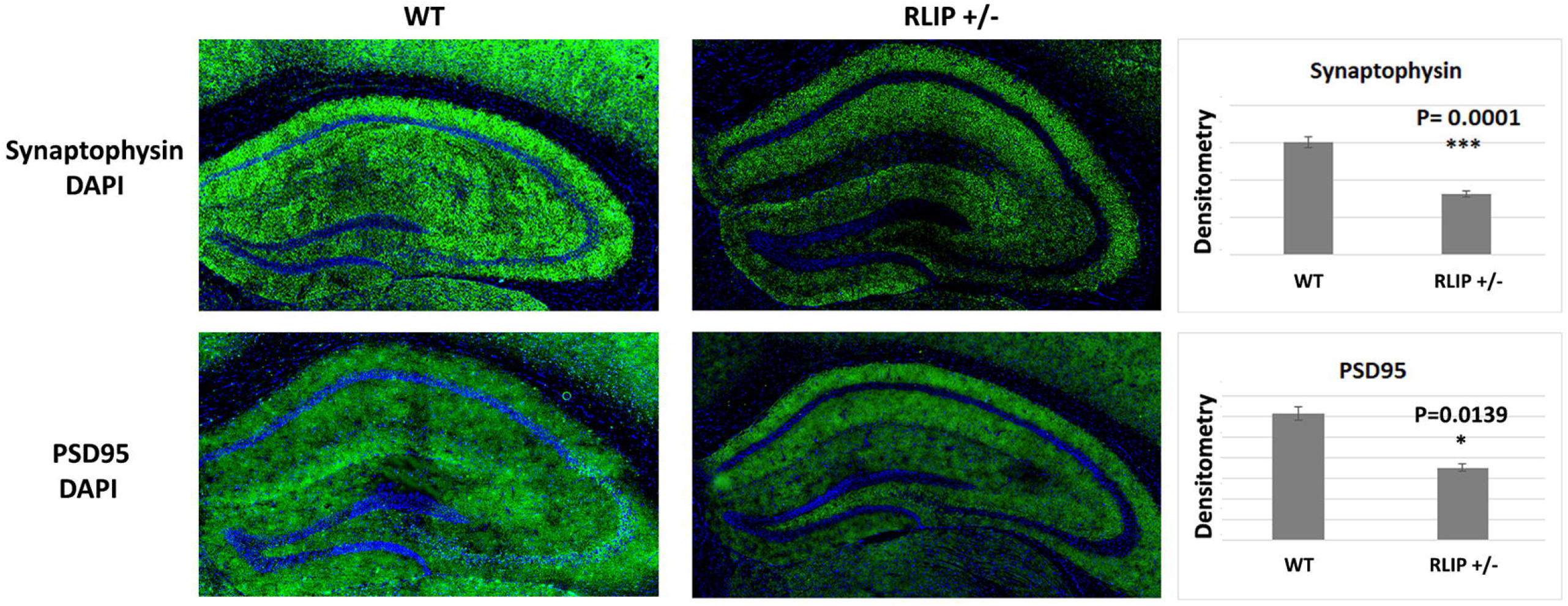
Immunofluorescence analysis of mitochondrial dynamics, biogenesis, synaptic proteins in the neurons of the hippocampus of WT and Rlip+/- mice. A comparison of mitochondrial dynamics, synaptic and biogenesis protein between Rlip+/+ and Rlip +/- mice showed that a significantly larger number of dentate gyrus neurons expressed Drp1, Fis1, PGC1a, Nrf1, synaptophysin and PSD95 in Rlip+/- compared with WT mice.

#### Mitochondrial biogenesis

**Figure 6B** indicates significantly increased immunoreactivities of mitochondrial biogenesis proteins (PGC1α, *p* = 0.0328 and Nrf1, *p*=0.043) in Rlip^+/-^ mice compared to WT mice. However, immunoreactivity of TFAM did not change in Rlip^+/-^ mice when compared to WT mice. These observations agree with our mRNA, immunoblotting data. Synaptic proteins: As shown in **Figure 6C**, synaptic proteins, synaptophysin (*p*=0.0001) and PSD95 (*p*=0.0139) were significantly reduced in Rlip^+/-^ mice, similar to published immunofluorescence data in APP and tau transgenic mouse models of AD (33, 34). It is worth noting that synaptophysin immunoreactivity reduced with high significance (*p*=0.0001).

### Rlip haploinsufficiency reduces OS and Nrf2-related antioxidant levels in mouse brain

In our previous studies, Rlip depletion induces OS in cells and several mouse tissues (35-38). Although the brain represents only 2% of body weight, it consumes 20% of the total body oxygen (refs). Therefore, increased OS in brain tissue would be expected to increase OS markers. We previously analyzed OS markers of wildtype (WT) and Rlip deficient mouse brain tissue. We demonstrated that LPO and total reactive aldehyde (TBARS, thiobarbituric reactive substances) were higher in the Rlip^+/-^ and Rlip^-/-^ than in WT, with a clearly significant sex-related difference, OS due to Rlip loss being greater in females (38-40). These data suggest the presence of OS in the brain from Rlip depleted mice. Since Rlip^+/-^ mice have high levels of OS. Therefore, we investigated effects on downstream NRF2-related antioxidant defenses. Results led to our original hypothesis that sensing of OS by NRF2 depends on normal Rlip protein expression.

Glutathione peroxidase is a mitochondrial antioxidant enzyme, scavenges free radicals produced in mitochondria, generally helps to maintain mitochondrial function, therefore, we measured GPx enzymatic activity in Rlip+/- and WT mice. As expected, GPx activity was significantly reduced in Rlip+/- mice relative to WT, indicating that mitochondrial function is defective in Rlip+/- mice **(Figure 7)**.

**Figure 7.**
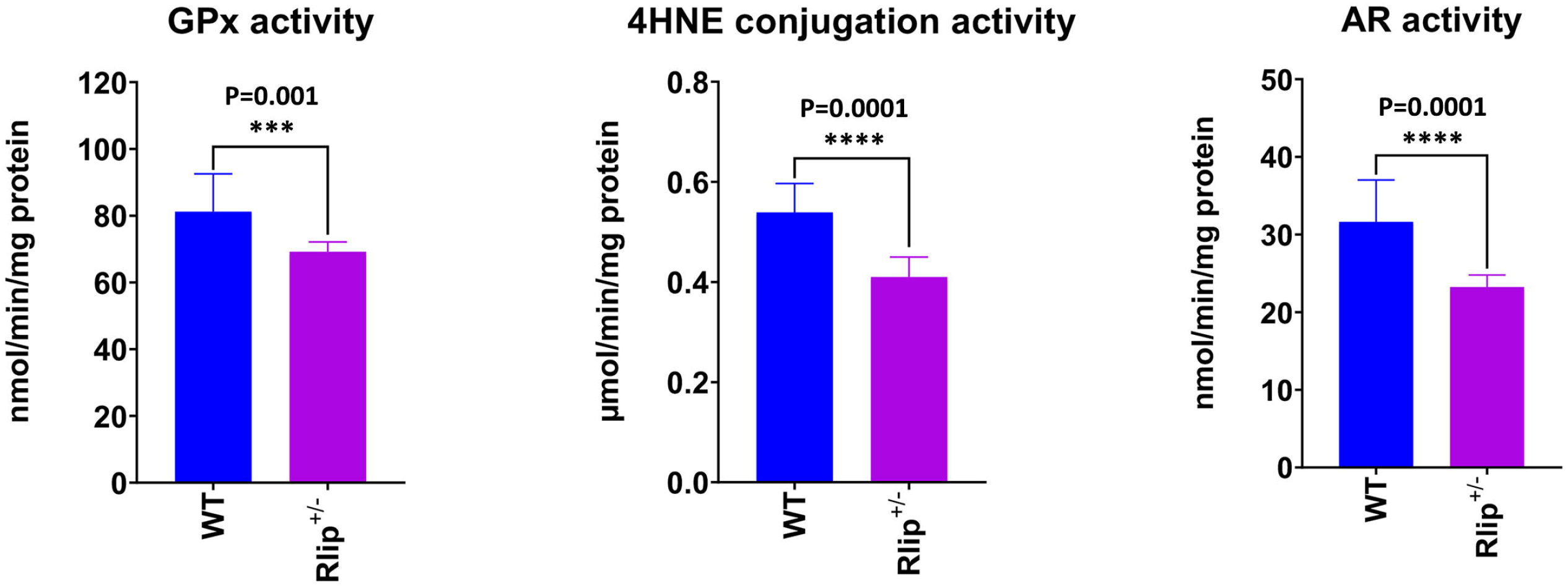
Antioxidant enzyme activities in Rlip^+/-^ mouse brains are lower than corresponding control wildtype mice. Brain tissue homogenates were prepared from three wildtype mice and three Rlip^+/-^ mice, and enzyme activity was measured in triplicate in individual homogenates as described in Methods. The activities were determined spectrophotometrically using standard methods established and validated in our lab. The difference in enzyme activity between WT and Rlip^+/-^ mice was significant for all tissues studied.

As shown in **Figure 7**, significantly reduced levels of aldose reductase in Rlip+/- mice (P=0.001) when compared to WT mice. Significantly reduced levels of 4-HNE conjugation activity were observed in Rlip+/- mice when compared with WT mice, indicating that lipid peroxidation is increased in Rlip+/- mice.

Our results confirmed lower activities of NRF2 regulated antioxidant enzymes aldose reductase (AR), glutathione S-transferase (GST) and glutathione peroxidase (GPx) in Rlip^+/-^ mice compared with age-matched wildtype mice (**Figure 7**). This finding is consistent with the results of many of our previous studies on sex-related differences in enzymes of xenobiotic metabolism such as GST and Rlip. The reduced activity of GST, GR, GPx, GGCS, GGT and G6PD in the brain and other tissues of Rlip knockout mice has been reported previously (35-38).

### Electron Micrographs Demonstrating Aberrant Mitochondria of Rlip^+/-^ mice

Mitochondria are primary energy sources. Additionally, OS associated mitochondrial dysfunction can be caused by changes in mitochondrial ultrastructure due to oxidative damage. To determine whether Rlip haploinsufficiency induces changes in mitochondrial ultrastructure, we compared mitochondrial morphology (number and length) in hippocampi and cerebral cortices from WT and Rlip^+/-^ mice using transmission electron microscopy. Consistent with previous studies (32, 41), electron microscopy revealed that Rlip^+/-^ mitochondrial number was significantly increased in the hippocampi (*p*=0.0023) and cortices (P=0.0055) from Rlip+/- mice, when compared with WT mice (**Figure 8**). The mitochondrial length was reduced in the hippocampi (*p*=0.0517) and cortices (P=0.0008). Overall, the differences in mitochondrial size, morphology and number were quite striking, similar to published reports in AD cell (42, 43)) and mouse models (33, 34). Therefore, Rlip^+/-^ mice are associated with OS-associated mitochondrial dysfunction, adding further validation to this knockout model.

**Figure 8.**
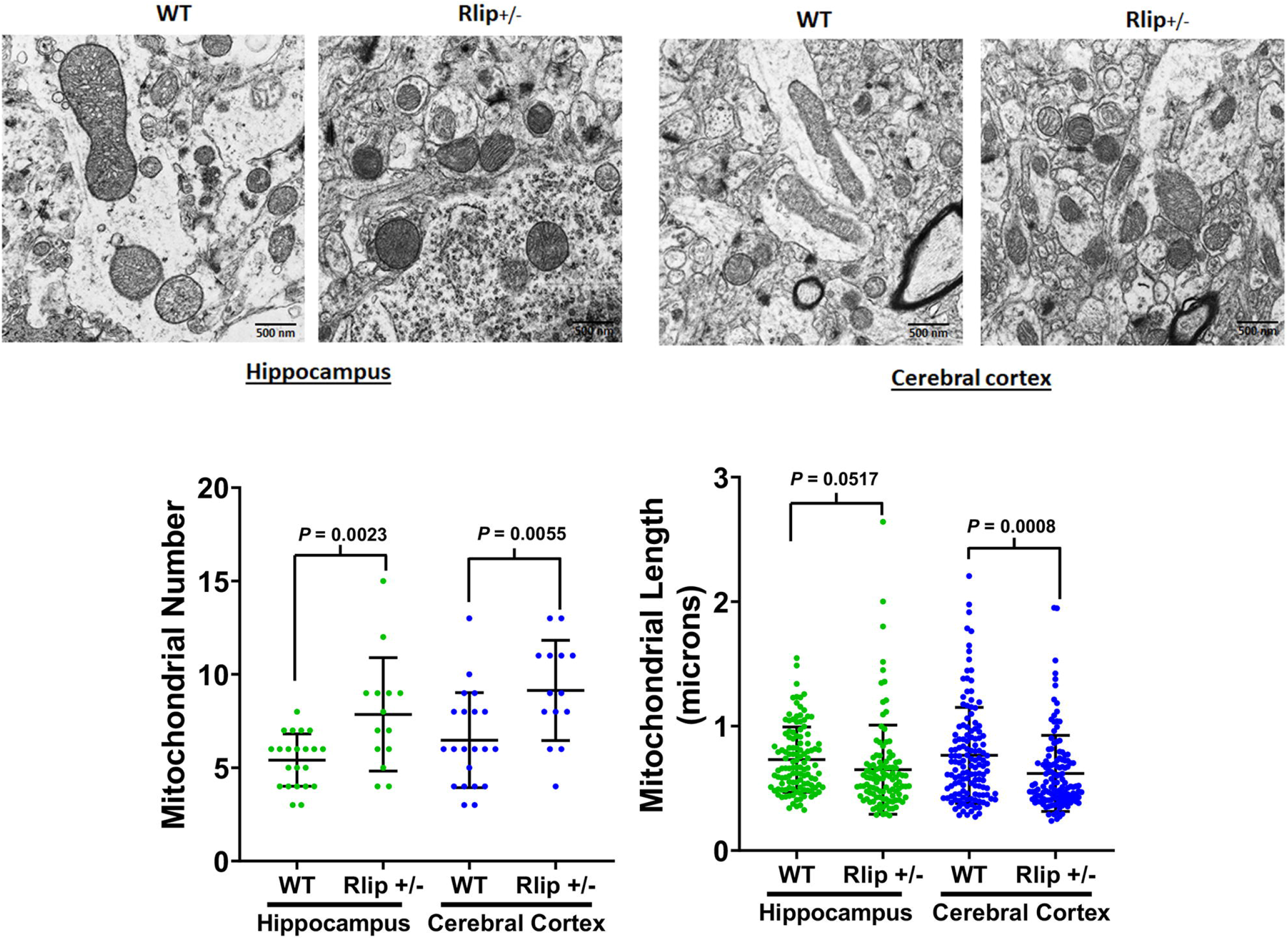
Electron micrographs demonstrating changes in mitochondria number and length in Rlip+/- mice relative to wildtype mice. Transmission electron microscopy was performed using hippocampal and cortical tissues from 10-month-old Rlip+/- (n=5) and wildtype mice (n=5). Mitochondrial number and length were assessed according to lab methods published in Kandimalla et al., 2018 (34) and Reddy et al., 2021 (42).

### Overlap in gene networks of AD, aging and inhibition of stress-activated gene expression in Rlip+/-

Rlip deficiency has global effects on gene expression and is associated with global promoter CpG island methylation abnormalities in liver tissue (31). We performed RNA-Seq transcriptomic analysis (n=3 Rlip^+/-^ and n=6 wildtype mouse brain and liver, with technical replicates) and analyzed the data by Ingenuity Pathway Analysis (IPA) software for effects on canonical pathways and upstream regulators. The results for differentially expressed canonical pathways (**Figure 9**) included a remarkable number of significantly affected pathways involved in neuronal diseases, most importantly synaptogenesis, NRF2 signaling, sirtuin signaling, and mitochondrial dysfunction, in both brain and liver tissues (**Figure 9A**). When the z-scores of differences were compared, we again found the changes to be significant with respect to synaptogenesis and NRF2-mediated oxidative stress.

**Figure 9.**
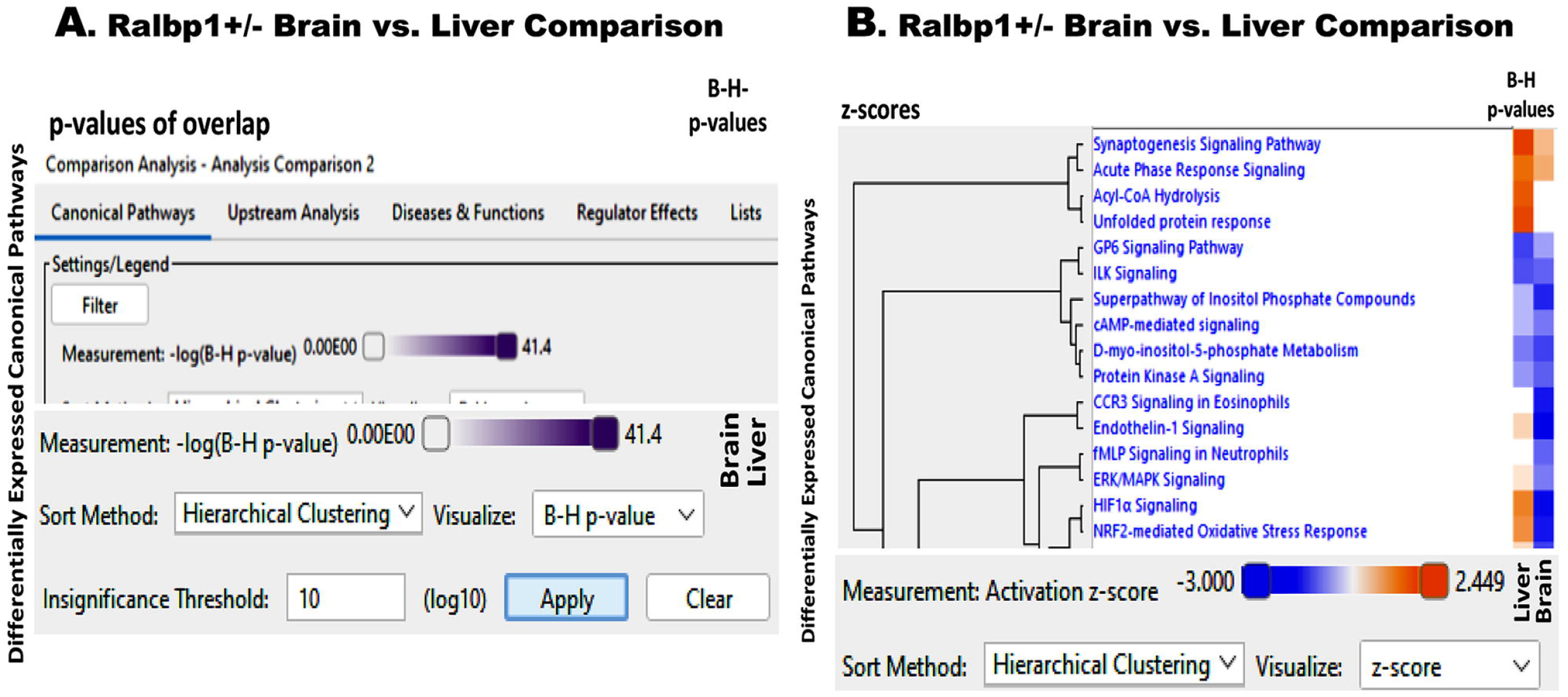
Gene Expression comparison between Ralbp1_HET brain versus liver by RNA-Seq: Whole genome transcriptomic analysis was performed by RNA-Seq on biological replicates for each group using an Illumina MiniSeq, and data were analyzed in the mouse mm10 assembly using NCBI RefSeq GRCm38.p1 gene annotations. Differentially expressed canonical pathways obtained using Ingenuity Pathway Analysis (IPA) were analyzed by Hierarchical Clustering methods using **(A)** p-value and **(B)** z-score methods. Validation of RNA-Seq results was performed by quantifying internal controls.

Perhaps most remarkably, the direction of change in NRF2-linked pathways was opposite between liver and brain, reduced in the latter and increased in the former (**Figure 9B**). This is a highly significant observation and shows that loss of Rlip has an inherently different effect on the brain as compared with the liver: the former would reduce the defense mechanisms in the brain while the latter would increase defense mechanisms in the liver. It is very important to investigate the underlying reasons for this remarkable difference. The upstream regulators most significantly affected (inhibited) were CREM, CRB1 CREBBP, CEBPA, NMDAR, GABRA1, and PSEN (Bias-corrected p values 1.15 e-7, 3.04e-7, 2.25e-5, 1.7e-4, 1.07e-6, 1.72e-9, 4.3e-4, 3.3e-3, respectively). Perhaps most relevant was APP (1.06e-6), the precursor protein of β-amyloid peptide (Aβ). All of these are strongly implicated in AD. As expected, based on the hypoglycemia and cancer resistance phenotypes of Rlip knockout mice, insulin and TP53 were included (p6.42e-4, 7.43e-4). Consistent with the known epigenetic effects of Rlip deficiency, DNMT3B was included (3.28e-3). The most relevant GSH utilization enzyme was LTC4, which is a druggable pathway in AD (4.57e-3). Pathways for CNS-active drugs/neurotransmitters included levodopa, dopamine, cocaine, methamphetamine, methylphenidate, thioridazine, and L-glutamate (p values ranging from 9.20e-3 to 1.26e-6). Consistent with the known functions of Rlip, estrogen signaling, mTOR, MAPK and HIF1α, unfolded protein (chaperone) response, clathrin dependent endocytosis, glutathione metabolism, oxidative stress and xenobiotic metabolism were among the top 25 pathways (p<1e-3).

### Transcriptional effect of Rlip Deficiency on Mitochondria

Preliminary evidence for the role of Rlip in neural synapses remains indirect. Though we did observe a reduction of synaptophysin expression, we have not yet visualized the neuronal synapses, studied clathrin-dependent endocytosis (CDE) in them or determined synaptic functions. Given the established and accepted function of Rlip in CDE it is likely that the proposed studies will experimentally confirm synaptic dysfunction. The transcriptomic results are showing widespread alterations in the CREB-signaling canonical pathway support this assertion and support a role of EP300 in the mechanisms through which inhibition of the normal physiological activation of Rlip by oxidative stress could exacerbate neuronal damage in AD (**Figures 10-12**).

**Figure 10.**
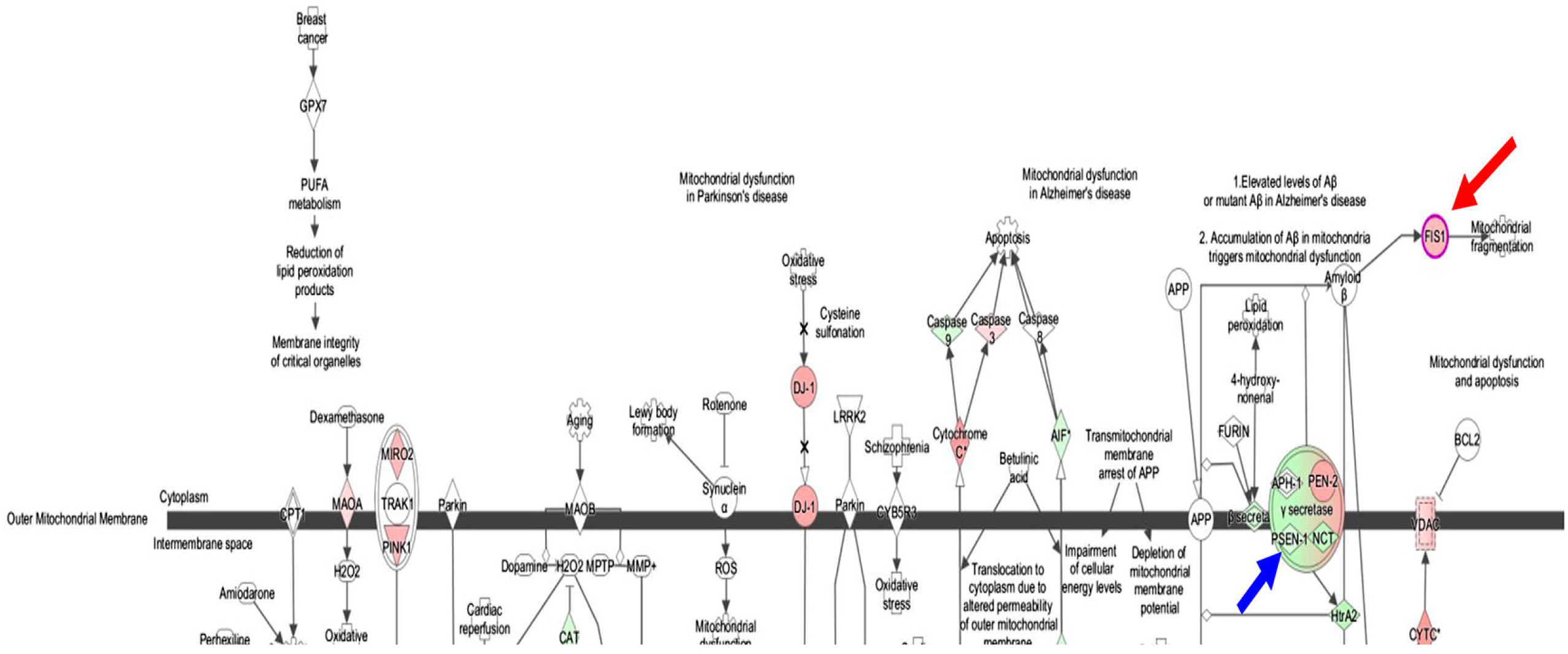
Transcriptional effect of Rlip deficiency on mitochondria. Brain RNA Seq data from Rlip^+/-^ mice (average of 3 males, 3 females, technical replicates) was overlaid on the canonical mitochondrial dysfunction pathway generated in IPA the arrows point out FIS1 (red), and PSN1 (blue).

**Figure 11.**
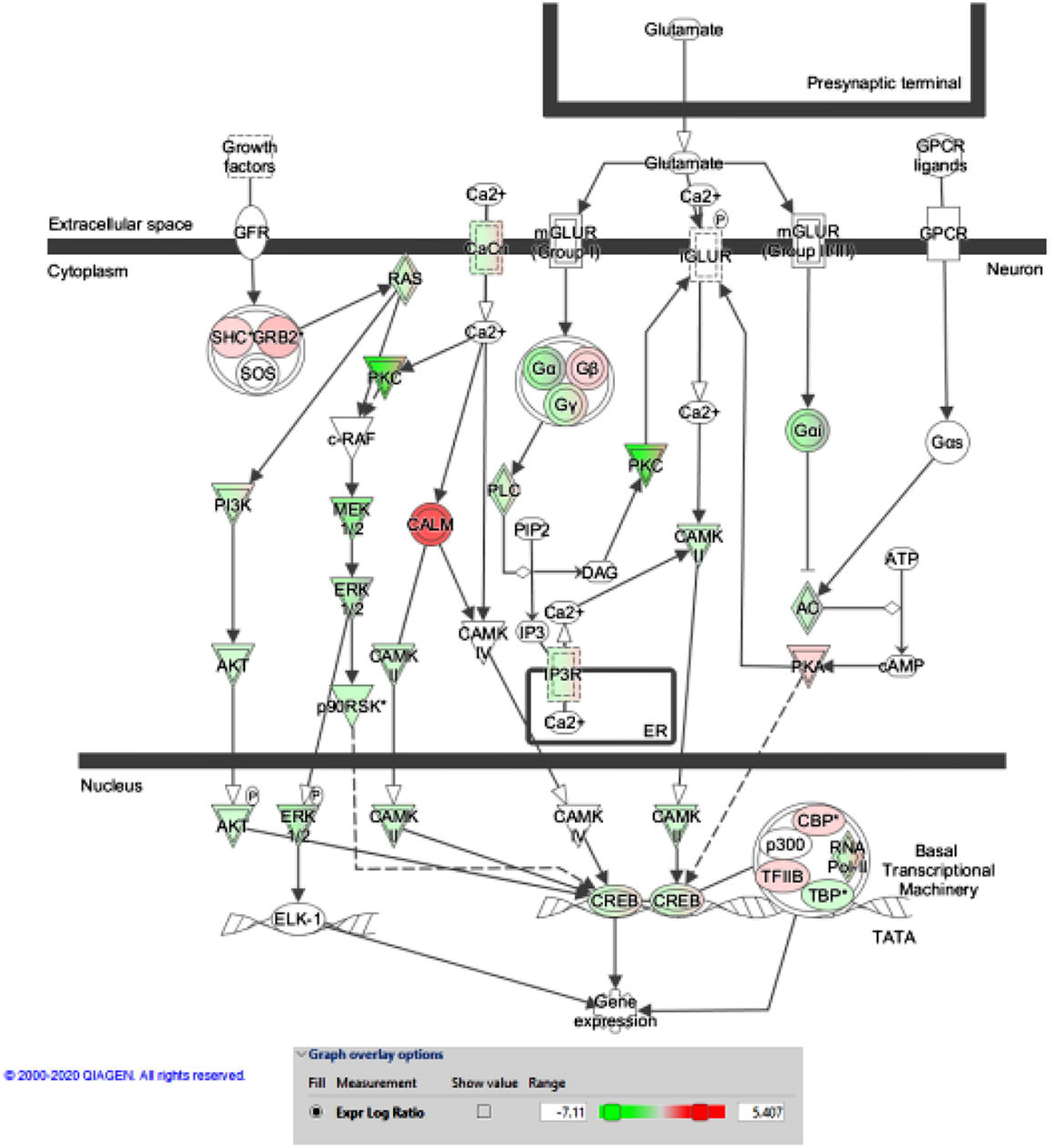
Transcriptomic effect of Rlip deficiency on the CREB signaling in neurons.

**Figure 12.**
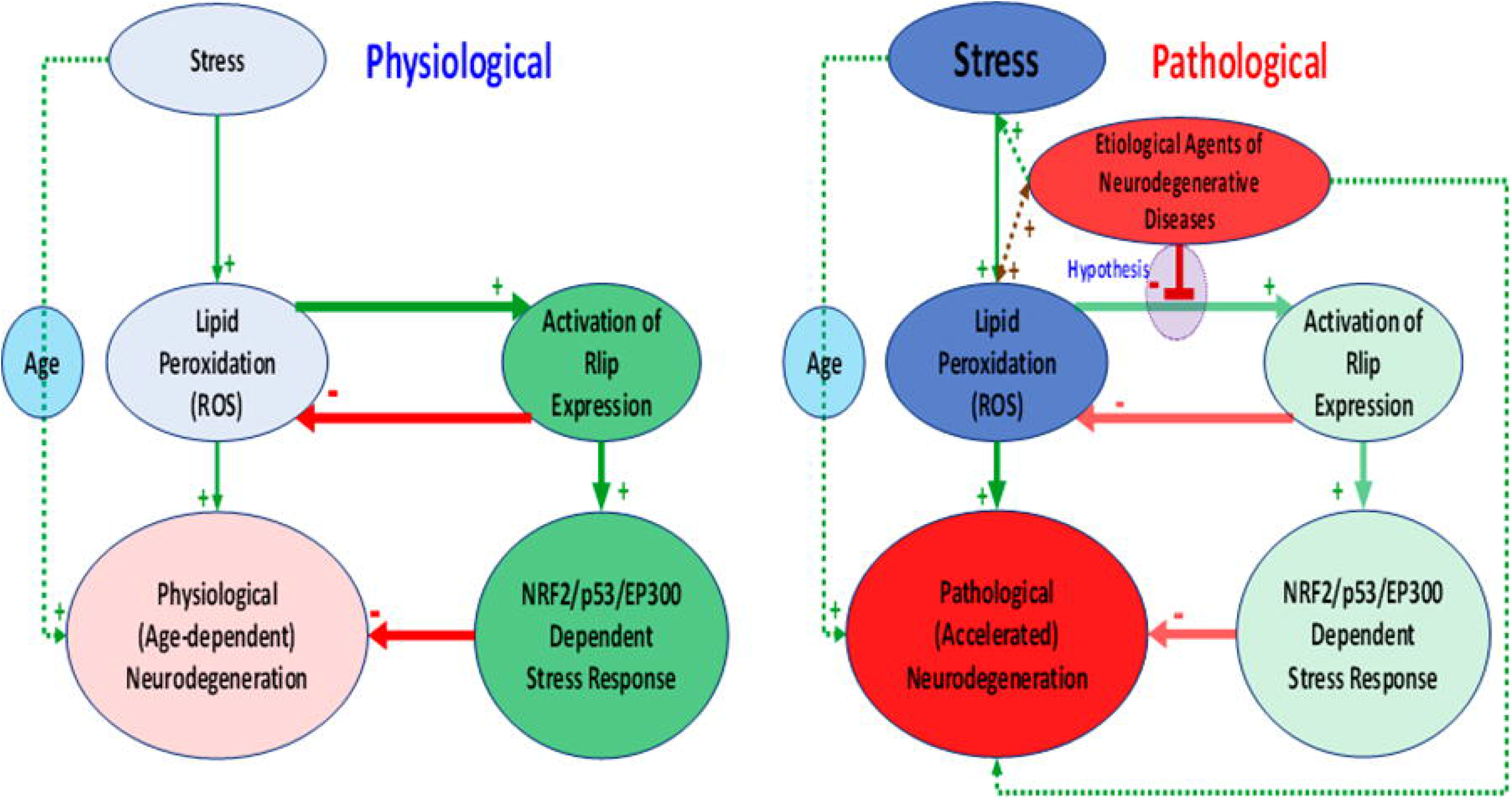
A hypothetical model of the role of Rlip-regulated expression of stress-defense genes controlled by NRF2, p53 and EP300. Red arrows represent inhibition and green activation. Note that according to this hypothesis, the etiological agents of neurodegenerative diseases inhibit activation of Rlip expression, which normally inhibits lipid peroxidation and neurodegeneration. Lighter shades indicate reduction.

## Discussion

The long-term goal of our study is to understand the impact of reduced expression of the Rlip gene in oxidative stress and mitochondrial dysfunction in Alzheimer’s disease progression and pathogenesis. In the current study, for the first time, we studied, Rlip heterozygous knockout mice (Rlip+/-) brain tissues for mRNA and protein levels of mitochondrial dynamics, biogenesis and synaptic genes, mitochondrial enzymatic activities, mitochondrial morphology and gene expression analysis. Our comprehensive and detailed analysis revealed that mitochondrial fission is increased, fusion & synaptic activities were reduced. Rlip+/- mice are cognitively deficits based on the open field, rotarod, Y-Maze and Morris Water Maze tests. Mitochondrial enzymatic activities, including Glutathione peroxidase, aldose reductase, 4-HNE conjugate activities were significantly reduced in Rlip+/- mice relative to age-matched wildtype mice. These features are similar to APP and Tau transgenic mouse models of Alzheimer’s disease. We studied about 7-12 months old Rlip+/- mice in comparison to wildtype mice (with similar genetic backgrounds). A limitation of our study findings is that one-time observation. Therefore, it is important to study our initial observations in a time-course manner, in other words, it is important to study several time-points starting with 2-, 6-, 12-, 18- and 24-month-old or until terminal stages of life.

### Cognitive behavior

The most predominant and striking sign in an AD patient is the progressive decline in cognition, primarily due to loss of neurons and synapses in the brain. In the current study, we used Morris water maze (MWM), Y-Maze, Open Field and Rotarod tasks to determine cognitive changes in Rlip^+/-^ mouse models. The open-field test is used for the measurements of general motor activity, exploratory behavior and measures of anxiety. Rlip^+/-^ mice showed reduced mobility, traveled less distance and their turn angle was significantly higher. We observed significant differences between Rlip^+/-^ and wildtype mice in overall activity levels or measures of anxiety. Rlip^+/-^ mice showed a significantly higher number of freezing episodes and increased freezing time. Our behavioral data suggest that Rlip^+/-^ mice displayed increased stress or anxiety. On the other hand, the general motor abilities of Rlip^+/-^ mice have been shown to be intact in the Morris water maze as swimming speed did not differ from wildtype animals.

The Rotarod Test suggests that Rlip^+/-^ mice had reduced motor coordination or fatigue resistance. Rlip^+/-^ mice displayed a phenotype of an AD mouse model with intact general motor activity and strength, but with impairments in motor coordination and balance present at the age of 10 months, most likely due to impaired cerebellar activity and increased anxiety. Spontaneous alternation, a measure of spatial working memory, is surprisingly slightly higher in Rlip^+/-^ mice. This suggests that Rlip^+/-^ mice tend to explore all three arms of the maze and have an innate curiosity to explore previously unvisited areas. Since Rlip^+/-^ mice show increased oxidative damage, mitochondrial dysfunction and reduced Nrf2 dependent enzyme levels, studying the behavior of these mice contribute to the understanding of the pathophysiology of neuropsychiatric disorders.

### Mitochondrial and synaptic gene expression and protein levels

Transcript levels of genes responsible for the expression of mitochondrial structural and synaptic genes in mouse brains were examined. Dentate gyrus results suggest increases in Drp1 and Fis1 and decrease in Mfn1, Mfn2, Synaptophysin and PSD95 mRNAs in homogenates similar to mutant HT22 cells as previously shown by Reddy *et*.*al* (44) and APP and Tau mouse models shown by Reddy Lab (33, 34) and others (45, 46). Through fusion and fission events, mitochondria maintain dynamics and bioenergetics to fine-tune the share of nutrients and electrochemical signals. Mitofusins Mfn1 and Mfn2 regulate mitochondrial fusion. Mfn1 plays a more prominent role in mitochondrial fusion, whereas Mfn2 mainly affects mitochondrial metabolism. On the other hand, fission is mediated by Drp1, interacts with another adaptor, Fis1, is known to be increased in AD patients. Increased overexpression of FIS1 protein in dentate gyrus neurons of Rlip ^+/-^ mice confirms our finding of qRT-PCR analysis. Reduced mRNA levels of synaptic genes in Rlip^+/-^ mouse brain are possibly responsible for synaptic damage and thus cognitive dysfunctions in Rlip deficient mice. Taken together, mRNA analysis results suggest that Rlip deficiency causes dysregulation of mitochondrial dynamics, and synaptic health. These results suggest that dysfunctional mitochondria and synaptic plasticity lead to cognitive impairment in these mice.

### Oxidative stress

Free radical scavenging antioxidants and glutathione (GSH)-mediated metabolism of reactive oxygen species are defenses against neurodegeneration that are impaired in Alzheimer disease and understanding their regulatory mechanism is of critical importance in devising novel therapies. We have previously observed that Rlip depletion downregulates Nrf2, and increases oxidative stress in cells. Severe impairment of Clathrin-dependent endocytosis (CDE) in Rlip knockout mice and lack of any studies on the effect of Rlip loss led to the idea that perhaps these mice have neurocognitive deficits due to defects in CDE as well as an enhanced rate of oxidative damage that is a characteristic of AD.

### Mitochondrial enzymes

In the current study, we report for the first time that Rlip depletion associated dysregulation of Nrf2 causes decreased levels of antioxidant/anti-electrophile enzymes, increased cognitive impairment and abnormalities of mitochondrial structure and synaptic proteins. Additional preliminary studies show that Rlip depletion epigenetically regulates several AD-linked genes including CREBBP and other genes implicated in neurocognition. We also found that synaptic neurotransmission, oxidative stress, CDE and retinoic acid signaling pathways were chief among those affected by promoter methylomic and transcriptomic alterations in Rlip knockout mice.

The postulated role of Rlip-depletion in enhancing the effects of oxidative stress in the brain raises the question of the mechanism of the damaging effect. Rlip depletion registered decreased activity of glutathione peroxidase activity for lipid hydroperoxides which is similar to that seen in the transgenic APP and tau mouse models of AD (33,34).

While the reduced glutathione peroxidase activity may be physiologically significant, most striking is Rlip^+/-^ mouse ability to conjugate 4-HNE. We also report here reduced activity of 4-HNE conjugating activity in Rlip^+/-^ mouse brain. In mammals, 4-HNE is the product of non-enzymatic degradation of oxidized polyunsaturated fatty acids, primarily of the abundant arachidonic acid. Moreover, 4-hydroxyalkenals of chain lengths different from the nine carbon atoms of 4-HNE may arise from peroxidation of other polyunsaturated fatty acids. As lipid peroxidation and the subsequent decomposition of the peroxides are thought to be non-enzymatic processes, 4-HNE and/or similar 4-hydroxyalkenals are expected to be generated in AD brain. Lower activity of aldose reductase, a key enzyme in the polyol pathway, was found in Rlip^+/-^ mouse brains. AR catalyzes nicotinamide adenosine dinucleotide phosphate-dependent reduction of glucose to sorbitol, leading to excessive accumulation of intracellular reactive oxygen species in AD brain. Thus, reduced detoxification of 4-HNE and other oxidants may be a major physiological role of Rlip depletion that leads to dementia in these mice.

### Mitochondrial morphology

Mitochondrial dysfunction is an important part of AD. These energy powerhouses are capable of self-replication but become partially dysfunctional in brain with AD through a variety of mechanisms, including a continuous vicious cycle of oxidative/electrophilic stress. To further establish the role of Rlip in mitochondrial function and morphology, we assessed mitochondrial size, morphology in the hippocampus and cortical tissues from WT and Rlip^+/-^ mice. Results suggest that mitochondrial size, morphology and number are quite strikingly different in Rlip^+/-^ mice.

### RNA-Seq Data

Our RNA-seq data demonstrate overlap in gene networks of AD, aging and inhibition of stress-activated gene expression in Rlip deficient mice. Because most of these genes are stress-induced, elevated oxidative stress in Rlip knockout mice should have caused upregulation; the experimental observation showed the opposite. Thus, we offer a provocative explanation: Rlip deficiency impairs the normal stress-induced transcriptional activation of many genes associated with AD. This is based on indirect evidence, and leads to a hypothesis that merits further examination because the APP mutant AD mice also have defective NRF2 regulation that is corrected by a compound that reduces AD pathology in cell and mouse models.

In summary, our initial characterization of Rlip+/- mice revealed several aspects of oxidative stress and mitochondrial and synaptic deficits, similar to transgenic APP and Tau mouse models of AD (33,34). Our initial observations are exciting, further warrant time-course analysis of Rlip+/- mice. If a partial deficiency of the Rlip gene, causing oxidative stress, mitochondrial and synaptic activities, it is worth overexpressing the Rlip gene in wildtype, meaning making transgenic Rlip mice and study overexpressed Rlip for cognitive behavior, oxidative stress, mitochondrial and synaptic activities. Alternatively, overexpress Rlip gene in APP and Tau mice and study, if overexpressed Rlip can reduce cognitive dysfunction and mitochondrial and synaptic deficits found in AD.

## Methods

### Euthanasia and Necropsy

Mice were used under Protocol # 18015 approved by the TTUHSC Institutional Animal Care and Use Committee. Brains were extracted, hemisected and, weighed. One hemisphere was fixed in formalin while another hemisphere was snap frozen at -80°C or homogenized fresh for molecular and biochemical analysis.

### Behavioral Testing

#### Open field maze

The open field maze evaluates locomotion but also anxiety because mice that are anxious will tend to avoid the center of the field. Each mouse was acclimated to the testing room for 60 min. They have then placed into the center of a dimly lit (20-30 lux) open-field apparatus (44 × 44 × 30 cm) and allowed to explore the entirety of the box for 10 minutes. In order to remove olfactory cues of urine from the surface of the apparatus, the apparatus was cleaned thoroughly with 30% ethanol between experiments. Noldus Ethovision software tracked the mouse’s movement in the OFM arena.

#### Morris Water Maze (MWM)

The Morris Water Maze test is specifically utilized for the measurement of hippocampal function and examining brain deficits in diseased mice (47). The test uses a tank filled with milky water to motivate the animal to escape onto the hidden platform just below the water’s surface. If the mouse reaches the platform before the allotted time, the test is terminated. Mice were trained and tested in a large, circular, galvanized steel pool (160 cm in diameter, 62 cm high, filled with 26°C ±1°C water to a height of 24 cm) to find a hidden platform (10 cm in diameter, located 1.5 cm below the surface of the water). A total of 10 male 10-month-old mice (5 WT and 5 Rlip^+/-^) were placed in a galvanized circular tank with cloudy water. The test was run for a maximum time of 60 seconds 4 times a day for 4 consecutive days. The AnyMaze automated tracking system (Stoelting Co., Wood Dale, IL) analyzed several dependent variables, including latency to escape, speed, path length, percent time spent in quadrants(s), number of platform crossings, the average distance from platform location, and percent thigmotaxia (tendency to swim around the edge).

#### Y-Maze

This is a working memory test used to measure the inclination for mice to explore new environments, revealing cognitive deficits in diseased mice (48). Typically, mice investigate a new arm of the maze in preference to returning to a previously visited arm. This involves many parts of a rodent’s brain including the prefrontal cortex, hippocampus, basal forebrain, and septum (48). Each mouse was placed in a Y-shaped maze with three grey plastic arms placed 120 degrees away from each other. The mice were initially placed at the end of one arm and allowed to freely explore the maze for five minutes. The camera above tracked the number of arm entries and trials in order to calculate the percentage of alternation using the AnyMaze software system. Normal mice remember the previous arm and naturally alternate between arms. An additional variation of the Y maze involves free alternation, in which the mouse is put in the Y maze for up to 15 minutes. Deviations from free alternation are logged as errors and indicate impairment of working memory. The Y maze was cleaned with MB-10 (Quip Laboratories Inc, Wilmington, DE 19802) after each trial. These calculations allowed us to quantify working memory deficits within these mice.

#### Rotarod test

Motor coordination and learning were assessed using the rotarod test. Mice were placed on an Ajanta three-compartment rotarod, with a horizontal rod rotating on its long axis that required mice to walk forward to avoid falling off of the rod (49). The mice were placed on the rod rotating at 18 RPMs, timed, and given a maximum running time of five minutes. This test was completed once a day for four consecutive days.

### RNA-Seq workflow and gene network analysis

RNA samples from WT and Rlip^+/-^ mouse liver and brain were prepared using the RNeasy mini kit from Qiagen (Valencia, CA) according to manufacturer instructions. RNA quality was assessed by microfluidic capillary electrophoresis using an Agilent 2100 Bioanalyzer. The RNA 6000 Nano Chip kit from Agilent Technologies (Santa Clara, CA) was used for subsequent library preparation. Sequencing libraries were prepared with the TruSeq RNA Sample Prep Kit V2 from Illumina (San Diego, CA) according to the manufacturer’s protocol. To satisfy the criteria for differential expression, a *p* value ≤ 0.01, fold change ≥ 2, and RPKM ≥1 in at least 2 samples was required.

### Quantification of mRNA expression using real-time RT-PCR

RNA-Seq results were confirmed using real-time quantitative PCR. cDNA using gene primers was performed on an ABI-7500 fast real-time PCR system using the SYBR Green master mix. 1 μg of total RNA from the liver and brain was used to synthesize cDNA by reverse transcription using the RT kit (Applied Biosystems). Total RNA was isolated from mouse brain using TRIzol™ Reagent (Thermofisher). Primers for qRT-PCR were designed using Primer Express Software (Applied Biosystems) for the housekeeping and test genes. Primer sequences are listed in Table 1. cDNA synthesis and qRT-PCR reactions were performed as described previously by Reddy *et*.*al*. (44) in brief, real-time polymerase chain reaction (qRT-PCR) was performed in replicate on a 7900HT Fast Real-Time PCR System (Life Technologies Corporation, Grand Island, New York, NY, USA) with PowerUp SYBR Green Master Mix (Thermo Fisher) in a total reaction volume of 20 μL containing 0.3 μM gene-specific primers. The cycling protocol was an initial denaturation at 95 °C, followed by 40 cycles of denaturation at 95 °C and annealing/extension at 60 °C. The bet-actin transcript was used as a reference for the normalization of mRNA levels. Gene expression levels were normalized for each individual animal by the ΔΔCt method (50).

**Table 1.**
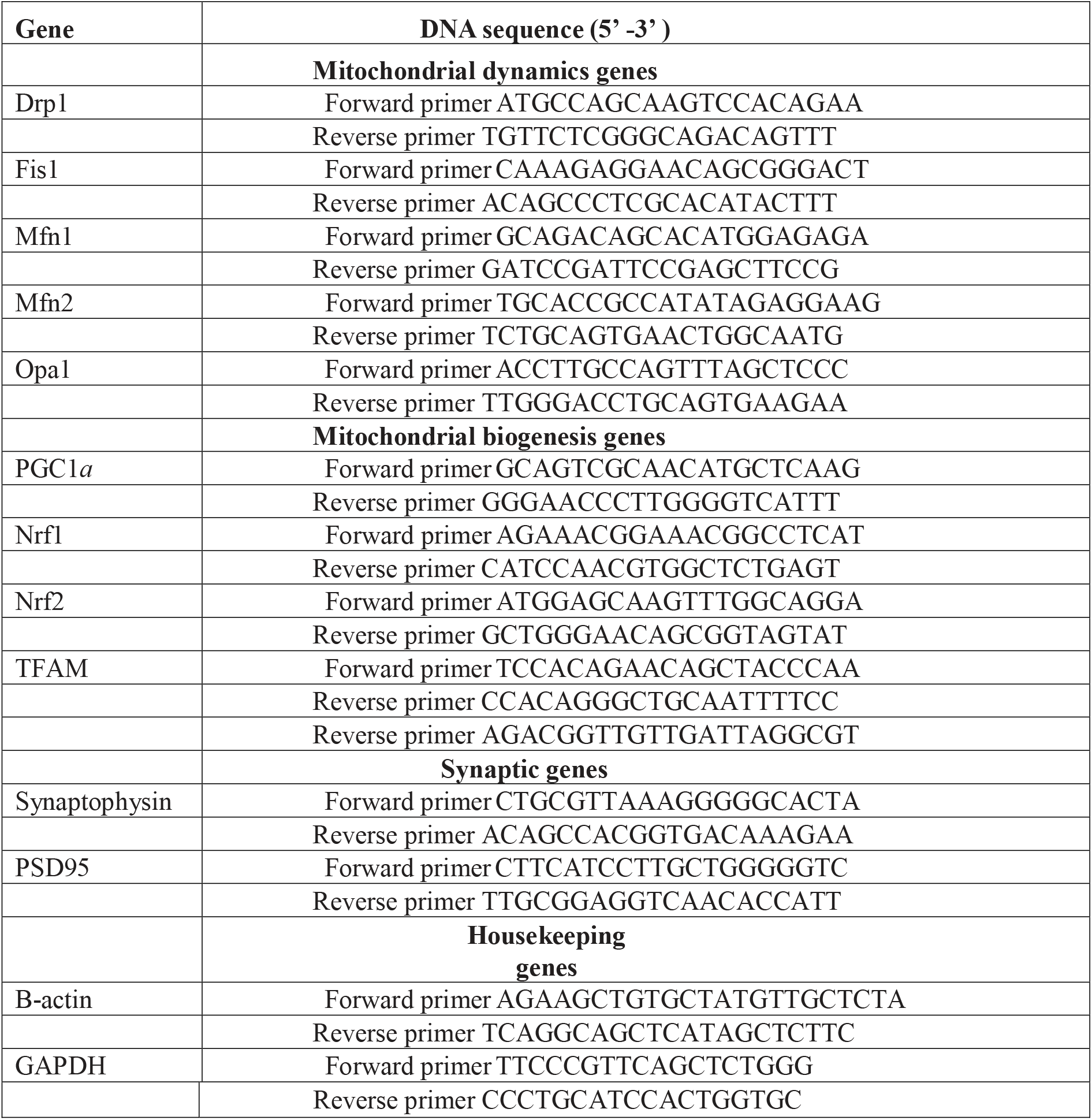
Summary of qRT-PCR oligonucleotide primers used in measuring mRNA expressions in mitochondrial dynamics, mitochondrial biogenesis, and synaptic, genes.

### Western blot analyses

Western blot analysis was performed using protein lysates prepared from Rlip+/- and wildtype mice. To outline the levels of mitochondrial biogenesis, dynamics and synaptic proteins, we have used beta-actin as an internal control. Details of antibody dilutions are given in Table 2. After treatment cells were lysed in 50μl cold RIPA lysis buffer (Millipore Sigma Aldrich Corporation, 20–188) for 60 mins on ice (vortex every 15 min interval) and centrifuged at 12,000 g for 11 min. After centrifugation, the supernatant was collected, and protein concentration was measured. 40 μg proteins were loaded and separated by SDS-PAGE gels (10%) electrophoretically and transferred to polyvinylidene difluoride membrane (Bio-Rad Incorporation, 10026933). Blocking was performed by adding 5% BSA for 60 min at room temperature on the shaker. After washing 2 times, primary antibody was added to the membranes overnight at 4-degree temperature. The membrane was washed 3 times with TBST and incubated with HRP (horseradish peroxidase)-labeled secondary antibodies for 1 hr at room temperature. Proteins were detected with chemiluminescence reagents (ECL, Thermo scientific, WA317048), and the band exposures were kept within the linear range. Bands from immunoblots were quantified using densitometry on a Kodak Scanner (ID Image Analysis Software, Kodak Digital Science, Kennesaw, GA). ImageJ software was used to quantify band intensity and determine statistical significance.

**Table 2.** Summary of antibody dilutions used in the immunoblotting analysis of mitochondrial dynamics, mitochondrial biogenesis, and synaptic proteins.

| **Marker** | **Primary antibody-species and dilution** | **Purchased from company, city and state** | **Secondary antibody, dilution** |
| --- | --- | --- | --- |
| DRP1 | Rabbit Polyclonal 1:1000 | Protein Tech Group | Donkey anti-rabbit HRP 1:10000 |
| FIS1 | Rabbit Polyclonal 1:500 | Novus Biologicals | Donkey anti-rabbit HRP 1:10000 |
| MFN1 | Mouse Monoclonal 1:1000 | Abcam | Donkey anti-mouse HRP 1:10000 |
| MFN2 | Mouse Monoclonal 1:1000 | Abcam | Donkey anti-mouse HRP 1:10000 |
| OPA1 | Rabbit Polyclonal 1:1000 | Novus Biologicals | Donkey anti-rabbit HRP 1:10000 |
| PGC1α | Rabbit Polyclonal 1:1000 | Novus Biologicals | Donkey anti-rabbit HRP 1:10000 |
| NRF1 | Rabbit Polyclonal 1:1000 | Cell Signaling Technology | Donkey anti-rabbit HRP 1:10000 |
| NRF2 | Rabbit Polyclonal 1:1000 | Novus Biologicals | Donkey anti-rabbit HRP 1:10000 |
| TFAM | Rabbit Polyclonal 1:2000 | Abcam | Donkey anti-rabbit HRP 1:10000 |
| PSD95 | Rabbit Polyclonal 1:1000 | Cell Signaling Technology | Donkey anti-rabbit HRP 1:10000 |
| SYNAPTOPHYSIN | Rabbit Polyclonal 1:3000 | Novus Biologicals | Donkey anti-rabbit HRP 1:10000 |
| B-Actin | Mouse Monoclonal 1:2000 | Millipore Sigma | Sheep anti-mouse HRP 1:10000 |

### Immunofluorescence analysis and quantification

Immunofluorescence analysis was performed using coronal hippocampal sections from WT and the Rlip^+/-^ mice on dynamics, biogenesis and synaptic proteins. Antibody dilutions of primary and secondary antibodies are given in Table 3. The sections (around 3–4 μm) were allowed to reach room temperature (∼30min) and rinsed at room temperature in PBS for 5 min. Then the sections were fixed in freshly prepared 4% paraformaldehyde in PBS for 10 min, washed with PBS and permeabilized with 0.1% Triton-X100 in PBS. They were blocked with a 1% blocking solution for 1 h at room temperature. All sections were incubated overnight with primary antibodies. After incubation, the sections were washed three times with PBS, for 5 min each. The sections were incubated with a secondary antibody conjugated with Alexafluor 488 (Invitrogen) for 1 h at room temperature. The sections were washed three times with PBS. After washing the sections, the coverslips were mounted on glass with Prolong Diamond antifade mounting medium (Thermo Fisher Scientific) media with DAPI for the identification of nuclei. Photographs were taken with an epifluorescence Nikon microscope system (Nikon Eclipse E600). To quantify the immunoreactivity of mitochondrial and synaptic antibodies for each treatment, 10-15 photographs were taken at 40X magnification as described in our previous publications (51, 52).

**Table 3.** Summary of antibody dilutions used in the immunofluorescence analysis of mitochondrial dynamics, mitochondrial biogenesis, and synaptic proteins.

| **Marker** | **Primary antibody-species and dilution** | **Purchased from company, city and state** | **Secondary antibody, dilution** |
| --- | --- | --- | --- |
| DRP1 | Rabbit Polyclonal 1:100 | Novus Biologicals | Donkey anti-rabbit<br>Alexa Fluor 488<br>1:200 |
| Fis1 | Rabbit Polyclonal 1:100 | Novus Biologicals | Donkey anti-rabbit<br>Alexa Fluor 488<br>HRP 1:200 |
| Mfn1 | Rabbit Polyclonal 1:100 | Thermofisher Scientific | Donkey anti-rabbit<br>Alexa Fluor 488<br>HRP 1:200 |
| PGC1α | Rabbit Polyclonal 1:100 | Novus Biologicals | Donkey anti-rabbit<br>Alexa Fluor 647<br>HRP 1:200 |
| NRF1 | Rabbit Polyclonal 1:100 | AbCam | Donkey anti-rabbit<br>Alexa Fluor 488<br>HRP 1:200 |
| TFAM | Rabbit Polyclonal 1:100 | Novus Biologicals | Donkey anti-rabbit<br>Alexa Fluor 647<br>HRP 1:200 |
| PSD95 | Rabbit Polyclonal 1:300 | Cell Signaling Technology | Donkey anti-rabbit<br>Alexa Fluor 488<br>HRP 1:500 |
| Synaptophysin | Rabbit Polyclonal 1:400 | Novus Biologicals | Donkey anti-rabbit<br>Alexa Fluor 488<br>HRP 1:600 |

### Aldose reductase assay

Brain lysates were prepared in G1 buffer (potassium phosphate 20mM, pH 7.0; BME-1.4mM and EDTA 2mM) and AR activity was assayed according to the method described previously (53). Briefly, the assay mixture contained 50 μM potassium phosphate buffer (pH 6.2), 0.4 mM lithium sulfate, 5 μM 2-mercaptoethanol, 10 μM DL-glyceraldehyde, 0.1 μM NADPH, and enzyme preparation (10μg/assay). The assay mixture was incubated at 37 °C and initiated by the addition of NADPH at 37 °C. The change in the absorbance at 340 nm due to NADPH oxidation was a Spectra Max plus (Molecular Devices) spectrophotometer.

### GST activity with 4-hydroxynonenal

4-HNE conjugating activity with glutathione was determined by methods described previously (54, 55). Assay mixture contained 100mM Kpi buffer (pH6.5), 0.5mM GSH and 0.1mM 4-HNE. Blank was without protein but with all other components, including 4-HNE (added at time 0), since the rate of non-enzymatic reaction of 4-HNE with GSH is relatively high. Measurement was started immediately after adding 4-HNE, since the reaction rate is linear with time for a short time only.

### Glutathione peroxidase assay with cumene hydroperoxide

Intracellular Glutathione peroxidase enzyme activity was determined in brain homogenates using Glutathione Peroxidase Assay Kit from abcam (Colorimetric; ab102530) (56). Briefly, tissues were homogenized in a hypotonic lysis buffer and sonicated. Glutathione peroxidase (GPx) oxidizes GSH to produce GSSG as part of the reaction in which it reduces cumene hydroperoxide. Glutathione reductase (GR) then reduces the GSSG to produce GSH, and in the same reaction consumes NADPH. The decrease of NADPH (measured at OD=340 nm) is proportional to GPx activity.

### Electron Microscopy

To determine the effects of Rlip deficiency on the mitochondrial number and size, we performed transmission electron microscopy in hippocampal and cortical sections of 12-month-old Rlip+/- mice relative to age-matched WT mice. Animals were perfused using the standard perfusion method—after successful perfusion, skin was removed on top of head and take the brain out, and post-fixed the brain for 2–3 h and/or definitely, and cut the hippocampal and cortical sections for transmission electron microscopy. 1× 1 mm thickness fixed sections from hippocampi and cortices were used for further analysis. Cut sections were stained for 5 min in lead citrate. They were rinsed and post-stained for 30 min in uranyl acetate and then were rinsed again and dried. Electron microscopy was performed at 60 kV on a Philips Morgagni TEM equipped with a CCD, and images were collected at magnifications of 36,000. The numbers of mitochondria were counted and statistical significance was determined, using one-way ANOVA.

### RNA-Seq workflow and gene network analysis

RNA samples from WT and Rlip^+/-^ mouse brain and liver were used. Ribosomal RNA was removed from total RNA using the RiboZero kit from Illumina and the resulting RNA was ethanol precipitated before cDNA synthesis. First-strand cDNA synthesis was performed using DNA polymerase I and RNase H. cDNA was end repaired, and 3’ end adenylated. Universal adapters were ligated followed by 10 cycles of PCR using Illumina PCR Primer Cocktail and Phusion DNA polymerase from Illumina. Subsequent library purification with Agencourt AMPure XP beads was validated with Agilent Bioanalyzer 2100, and quantified with Life Technologies’ Qubit from ThermoFisher (Waltham, MA). Sequencing was conducted on Illumina Hiseq 2500 with single end 50 bp reads. Reads were aligned using Tophat v2.0 to mouse reference genome mm9. The expression level of Refseq genes was counted and normalized using the TMM method (31, 57) and differential expression analysis was conducted using a linear model based on negative binomial distribution using :edgeR:. RPKM (**r**eads **p**er **k**ilobases per **m**illion mapped reads) was defined as a number of reads / (gene length/1000 * a total number of reads/1,000,000). Analyses were performed by censoring the lowest expressed genes or using log_2,_(RPKM+0.1) expression levels. To satisfy the criteria for differential expression, a *p* value ≤ 0.01, fold change ≥ 2, and RPKM≥1 in at least 2 samples was required. RNA-Seq results were confirmed using real-time quantitative PCR. cDNA using gene primers was performed on an ABI-7500 fast real-time PCR system using the SYBR Green master mix. 1 μg of total RNA from the liver was used to synthesize cDNA by reverse transcription using the RT kit (Applied Biosystems).

## Declarations Ethics Approval

The presented research is in compliance with ethical standards.

## Consent to Participate

Not applicable.

## Consent to Publish

All authors agreed to publish the contents.

## Research involving Human Participants and/or Animals

This study was carried out in strict accordance with the recommendations of U.S. National Institutes of Health Guide for the Care and Use of Laboratory Animals. The Institutional Animal Care and Use Committee (IACUC approval #18015) approved the protocol. All procedures were performed under anesthesia and all efforts were made to minimize the pain and suffering of the animals.

